# Targeted DamID in *C. elegans* reveals a role for LIN-22 and NHR-25 in epidermal cell differentiation

**DOI:** 10.1101/2020.12.17.423252

**Authors:** Dimitris Katsanos, Michalis Barkoulas

## Abstract

Transcription factors are key players in gene networks controlling cell fate specification during development. In multicellular organisms, they can display complex patterns of expression and binding to their targets, which necessitates tissue-specific characterisation of transcription factor-target interactions. Here, we focus on *C. elegans* seam cell development, which is used as a model of robust epidermal stem cell patterning. Despite our knowledge of multiple transcription factors playing a role in epidermal development, the composition of the gene network underlying cell fate patterning remains largely unknown. We introduce Targeted DamID (TaDa) that allows tissue-specific transcription factor target identification in intact *C. elegans* animals without cell isolation. We employ this method to recover putative targets in the epidermis for two transcription factors, the HES1 homologue LIN-22 and the NR5A1/2 nuclear hormone receptor NHR-25. Using single-molecule FISH (smFISH), we validate TaDa predictions and reveal a role for these transcription factors in promoting cell differentiation, as well as an unusual link between a HES factor and the Wnt signalling pathway.

Our results expand our understanding of the epidermal gene network and highlight the power of TaDa to dissect the architecture of tissue-specific gene regulatory networks.

## Introduction

Development of animals and plants involves the reproducible formation of complex organisms starting from single cells. This remarkable transformation requires genetic information to be decoded in a manner that allows cell type diversity to emerge through spatiotemporal control of gene expression. In this process, transcription factors (TFs) are prominent players as they participate in gene regulatory networks that govern robust cell fate specification, often acting in a combinatorial way (Reilly et al., 2020). Therefore, elucidating the architecture of developmental gene networks by identifying the participating TFs, as well as their biologically relevant targets, is fundamental for increasing our mechanistic understanding of how specialised cells and tissues are formed (Davidson, 2010; Oliveri and Davidson, 2007).

An important type of cells in the *C. elegans* epidermis are the seam cells, which follow a stem cell-like pattern of symmetric and asymmetric divisions throughout larval development to produce most epidermal nuclei (Chisholm and Hsiao, 2012; Sulston and Horvitz, 1977). Proliferative symmetric divisions occur at the second larval stage and increase the stem cell pool, whereas asymmetric cell divisions at each larval stage generate differentiated hypodermal or neuronal cells, while maintaining the total seam cell number through self-renewal (Sulston and Horvitz, 1977). The highly reproducible nature of *C. elegans* development allows the use of this model to reveal the mechanisms underlying the balance between cell proliferation and differentiation (Joshi et al., 2010). As in other stem cell contexts, Wnt signalling plays a key role in seam cell maintenance and the regulation of seam cell division patterns (Gleason and Eisenmann, 2010). In the canonical Wnt pathway, β-catenin is targeted for degradation in the absence of Wnt ligand binding to receptors. Upon Wnt receptor activation, β-catenin is stabilised, enters the nucleus and, along with the TF POP-1/T-cell factor/lymphoid enhancer factor (TCF/LEF), activates Wnt target genes (Sawa and Korswagen, 2013). The Wnt/β-catenin asymmetry (WβA) is a modified version of the Wnt pathway adapted for the purposes of asymmetric cell division, where asymmetric distribution of Wnt pathway components polarises the mother cell and leads to asymmetric inheritance of the potential for Wnt pathway activation in the two daughter cells following division (Lam and Phillips, 2017).

Besides Wnt signalling, a number of conserved TFs have been identified to influence cell fate decisions in the epidermis. For example, RNT-1 is the *C. elegans* Runx homolog and has been shown to act together with its binding partner BRO-1/ CBFβ to promote symmetric seam cell divisions (Kagoshima et al., 2007; van der Horst et al., 2019). Mutations in Runx genes and CBFβ are known to cause various leukaemias in humans (Cameron and Neil, 2004), and mutations or overexpression in *C. elegans* lead to seam cell loss or tissue hyperplasia respectively (Nimmo et al., 2005). Another example is the HES1 homologue and basic helix-loop-helix (bHLH) TF LIN-22, which acts to suppress neurogenesis in seam cells (Wrischnik and Kenyon, 1997) and maintain correct patterning, possibly by antagonising the Wnt signalling pathway (Katsanos et al., 2017). The GATA family of TFs is linked to various types of cancer in humans (Zheng and Blobel, 2010), and related factors in *C. elegans*, such as ELT-1 and EGL-18, are thought to specify seam cell fate (Gorrepati et al., 2013; Koh and Rothman, 2001; Smith et al., 2005). Finally, TFs of the nuclear hormone receptor (NHR) family have been reported to be play a role in epidermal development (Antebi, 2006; Miyabayashi et al., 1999). A key example is the NR5A1/2 homologue NHR-25, which regulates the establishment of cell-to-cell contacts in the seam, while it acts more broadly in vulval development, molting and neurogenesis in the T lineage (Chen et al., 2004; Gissendanner and Sluder, 2000; Hajduskova et al., 2009; Hayes et al., 2006; Shao et al., 2013; Šilhánková et al., 2005).

Despite the gain in knowledge on individual TFs playing a role in epidermal development, we still have a limited understanding of the underlying gene network and how this drives the decision between cell proliferation and differentiation. This is largely due to the fact that most identified components are still disconnected to each other, or when interactions are known, these remain at the genetic level. Identification of direct TF targets has been predominantly pursued by ChIP experiments in *C. elegans* (Araya et al., 2014; Celniker et al., 2009; Kudron et al., 2018). DNA adenine methyltransferase identification (DamID) (van Steensel et al., 2001; van Steensel and Henikoff, 2000), offers an alternative way to reveal targets by fusing a TF of interest to the Dam methyltranferase from *E. coli*. This enzyme mediates site-specific methylation in the adenine of GATC sequences therefore it can leave methylation marks to the DNA wherever a TF binds (Schuster et al., 2010; van de Walle et al., 2020). A key requirement in DamID methods is keeping the levels of Dam expression very low to avoid toxicity and saturated methylation of DNA (Pindyurin et al., 2016; van Steensel and Henikoff, 2000). An additional challenge in multicellular systems like *C. elegans* is to characterise DNA-protein interactions with tissue-specific resolution. This has been previously achieved using recombinase-based systems to allow expression of Dam fusions at low levels defined by the basal transcription from non-induced heat-shock promoters (Gómez-Saldivar et al., 2020; Harr et al., 2020; Pindyurin et al., 2016). Alternatively, Targeted DamID (TaDa) allows tissue-specific target identification (Southall et al., 2013) and has been successfully employed in *Drosophila* to dissect mechanisms of neuronal fate determination (Sen et al., 2019; Vissers et al., 2018), as well as in mammalian stem cell lines to characterise the binding profile of pluripotency factors (Cheetham et al., 2018). Low levels of Dam-fusion expression are achieved in TaDa with a specific transgene configuration where an unrelated primary ORF (usually *mCherry)* is introduced followed by 2 STOP codons and a frameshift preceding the Dam-fusion as a second ORF (Figure 1A). Due to the universal property of eukaryotic ribosomes to reinitiate translation after a STOP codon at a reduced frequency (Kozak, 2001, 1987), this transgene design results in low levels of expression of the Dam-TF fusion in the tissue of interest as defined by the specific promoter used (Figure 1A).

**Figure 1:**
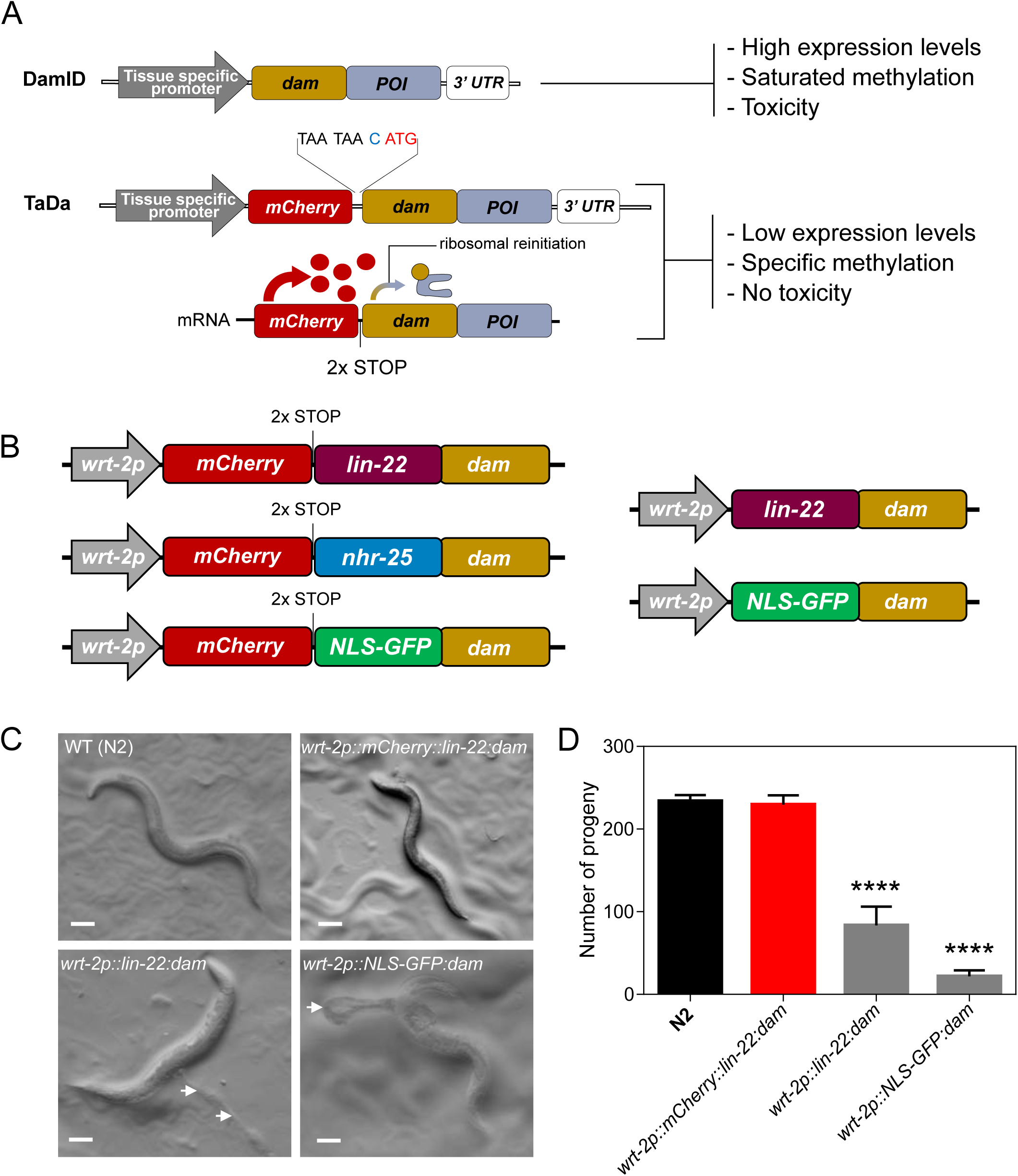
TaDa transgene design prevents animal toxicity. **(A)** Schematic showing the bicistronic design of TaDa with a primary ORF of *mCherry* followed by two STOP codons and a frameshift, which permits low levels of tissue-specific expression of a fusion between the protein of interest (POI) and Dam. **(B)** Illustration of the key features of single-copy transgenes used in this study for LIN-22 and NHR-25 target identification by TaDa (left). Transgenes used to assess the requirement of an *mCherry* primary ORF to prevent toxicity and high methylation levels are shown on the right. **(C)** Representative images of adult wild type (WT) and transgenic animals carrying TaDa fusions. Note that animals carrying *lin-22:dam* or *NLS-GFP:dam* fusions, that is in the absence of the primary *mCherry* ORF, show aberrant phenotypes. White arrows point to internal tissue outside the animal body. **(D)** Quantification of brood size in the above strains. Note defect in brood size in animals carrying *lin-22:dam* and *NLS-GFP:dam* fusions (n=15). Scale bars in C are 100 μm. In D error bars indicate standard error of the mean and black stars indicate statistically significant differences in the mean with a one-way ANOVA followed by a Dunnett’s test (* *p*<0.05, **** *p*<0.0001).

In this study, we demonstrate that TaDa is a powerful method to identify TF targets in *C. elegans*. We employ TaDa to find targets of LIN-22 and NHR-25 in the epidermis. Using single molecule FISH (smFISH), we validate changes in target expression upon perturbation of LIN-22 or NHR-25 activity. Our results suggest a role for these two TFs in promoting cell differentiation by repressing stem cell promoting factors. We also reveal a direct link between LIN-22 and the Wnt signalling pathway through the Frizzled receptor *lin-17*. Our findings expand our knowledge of the gene network underlying epidermal cell fate decisions and provide a methodological framework for resolving regulatory networks in specific nematode tissues.

## Results

### TaDa circumvents Dam-associated toxicity and allows the recovery of TF-specific methylation profiles

To enrich our understanding of the gene network underlying epidermal cell fate patterning, we chose to study two TFs, the bHLH factor LIN-22 and the nuclear hormone receptor NHR-25, which are both key players in epidermal development (Chen et al., 2004; Gissendanner and Sluder, 2000; Wrischnik and Kenyon, 1997). We decided to focus on LIN-22 because previous work implicated this factor in Wnt-dependent seam cell patterning, although its direct targets remained unknown (Katsanos et al., 2017; Wrischnik and Kenyon, 1997). On the other hand, NHR-25 targets had been previously studied using ChIP-seq (Araya et al., 2014; Shao et al., 2013), although direct targets in the epidermis were not known. To profile TF targets by TaDa, we constructed transgenes consisting of TF-Dam fusions under the *wrt-2* promoter, which drives expression primarily in the seam cells. We included a *C. elegans* optimised *mCherry* as the primary ORF, followed by 2 STOP codons and an extra nucleotide for frame-shift before the TF-Dam (*lin-22:dam* and *nhr-25:dam)* or control (*NLS-GFP:dam)* fusions (Figure 1B). We also designed a versatile backbone to allow the assembly of N- or C-terminal fusions of any other transcription factor with Dam and using any promoter of interest (Figure S1A).

Induced ubiquitous Dam expression by a heat-shock promoter has been shown to produce saturated methylation and toxicity (Schuster et al., 2010; van Steensel and Henikoff, 2000), but the effect of tissue-specific expression of Dam-fusions had not been previously investigated in *C. elegans*. To test this, we constructed *lin-22:dam* fusions with and without *mCherry* as the primary ORF (Figure 1B), and inserted these into the genome as single-copy transgenes. We found that transgenic lines lacking *mCherry* showed developmental defects, as well as significantly reduced brood size compared to animals containing the *mCherry* primary ORF, which appeared comparable to wild-type controls (Figure 1C, D). Aberrant phenotypes in the absence of the primary ORF were likely due to high levels of Dam expression because they were observed in strains carrying both *lin-22:dam* and *NLS-GFP:dam* fusions, and correlated with increased amount of DNA methylation (Figure S1B). We therefore conclude that the TaDa transgene configuration reduces toxicity in *C. elegans*, which is consistent with what has been reported in other systems (Southall et al., 2013; van Steensel and Henikoff, 2000).

We then tested the single-copy transgenics harbouring TaDa transgenes for expression and functionality of each fusion. The expression of *mCherry* was used as proxy to confirm the spatial localisation of the TF-Dam fusions. Microscopy at the L4 stage revealed low *mCherry* expression in the seam cells, as expected for single copy transgenes (Figure S2A). Despite the TaDa design that results in low level of expression of the TF-Dam fusions, their functionality can still be assessed using a multi-copy transgenesis assay, where an increase in copy number may allow to rescue loss-of-function mutations or induce ectopic phenotypes. For example, *lin-22(icb38)* mutants show an increase in the number of postdeirid (PDE) neurons, and this phenotype was suppressed in transgenics carrying the *lin-22:dam* construct as an extrachromosomal array (Figure S2B). Furthermore, the *nhr-25:dam* fusion was sufficient to disrupt seam cell patterning as a multi-copy array (Figure S2C).

We sequenced amplicons derived from methylated genomic sequences for both TFs at two developmental stages (L2 and L4) and processed the data using the damidseq pipeline (Marshall and Brand, 2015). We generated normalised aligned read count maps to evaluate replicate reproducibility and assess the level of correlation between samples. High level of reproducibility was observed for all fusions between biological replicates (Figure S3A, B) with higher Pearson correlation coefficients for comparisons within a given *TF:dam* fusion (Figure S3C). Importantly, control *NLS-GFP:dam* samples were found to cluster separately from TF-Dam samples using principal component analysis (Figure S3D), and separate clustering was observed for the *lin-22:dam* and *nhr-25:dam* samples (Figure S3E). These results highlight that TF-Dam fusions produce specific and distinct methylation patterns compared to control fusions.

### NHR-25 and LIN-22 binding profiles overlap with regulatory regions of the genome

To investigate the profile of signal enrichment across the genome, we compared *TF:dam* and *NLS-GFP:dam* sample pairs and calculated the normalised log_2_(*TF:dam/NLS-GFP:dam*) ratio scores per GATC fragment based on normalised aligned read count maps (an example covering 1Mb of sequence is shown in Figure S3F). To address whether the observed signal enrichment is likely to reflect TF binding, we focused on putative targets of LIN-22 and NHR-25 proposed by genetic analysis or ChIP-seq experiments respectively. For example, we observed significant enrichment in the *lin-22:dam* profiles near the Frizzled receptor *lin-17*, the Hox gene *mab-5* and the *C. elegans achaete-scute* homologue *lin-32*, which are reported to be genetic interactors of *lin-22* (Katsanos et al., 2017; Wrischnik and Kenyon, 1997) (Figure 2A). With regard to the *nhr-25:dam* profiles, significant enrichment was found in regions near the putative target genes *idh-1* and *rpl-3,* as well as the *nhr-25* locus itself, supporting previous evidence for self-regulation (Shao et al., 2013) (Figure 2B). Qualitative inspection of the enrichment profiles suggested a preference for enrichment in intergenic regions (Figure 2A, B). Indeed, aggregate genome-wide signal profiles showed increased average enrichment scores in upstream sequences directly proximal to the transcriptional start site (TSS) of protein coding genes (Figure 2C).

**Figure 2:**
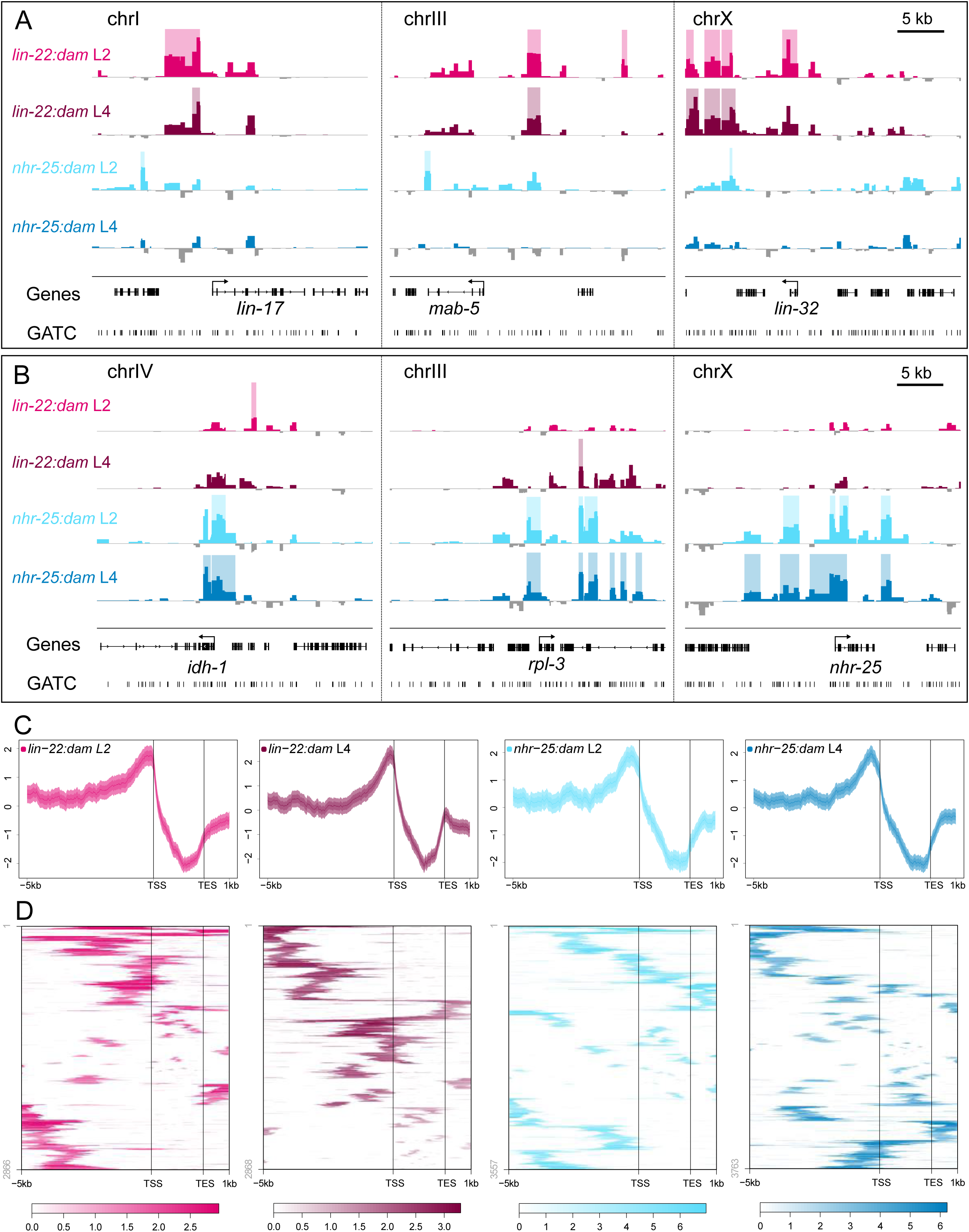
LIN-22 and NHR-25 binding signal is enriched in upstream gene regulatory regions. **(A-B)** Examples of signal profiles in selected regions associated with putative target genes. Shaded regions indicate statistically significant peaks (FDR<0.05). The Y-axes represent log_2_*(TF:dam*/*NLS-GFP:dam)* scores (Data range for *lin-22:dam*: -1 – 3.5, and for *nhr-25:dam*: -3 – 8). Scale bar length is 5 kb as indicated. **(C)** Aggregation plots depicting average enrichment scores in 10 bp bins for regions of equal length across all of the specified genomic features indicated on the X-axis. Strong enrichment preference is seen upstream to gene regions. Plots show 5 kb upstream of the TSS of genes to 1 kb downstream of the TES, with gene bodies pushed into a 2 kb pseudo-length. Y-axes are z-scores for the plotted sequence length and shaded areas represent 95% confidence intervals. **(D)** Heatmaps representing the hierarchically clustered localisation and enrichment score of all statistically significant peaks (FDR<0.05) within 5 kb upstream and 1 kb downstream of a gene.

Statistical processing of the signal profiles was used to identify significant peaks. We found 1965 and 1972 peaks for *lin-22:dam* at the L2 and L4 stage, and 2044 and 2169 peaks for *nhr-25:dam,* a complete list of which is shown in Table S1. Hierarchical clustering of the localisation and score of those peaks that lie between 5 kb upstream and 1kb downstream of genes, was consistent with a preference for signal enrichment in upstream to TSS regions (Figure 2D). The localisation of the peaks in relation to genes was further dissected by assigning peaks to their single closest gene. For all profiles, the majority of the peaks (>94%) were assigned to genes, with the largest proportion of peaks (between 41.2% and 47.3%) found to be upstream to or overlapping the TSS of their assigned gene (Figure 3A). From the peaks classified as upstream of genes, approximately half (46% - 54%) localised within the first 2 kb upstream. A significant proportion of peaks were found to be within genes, but around a quarter of those (24.3% - 28.5%) were exclusively residing within introns, which are known to contain regulatory elements in *C. elegans* (Fuxman Bass et al., 2014) (Figure 3A). In stark contrast, a very small percentage of peaks, (4.2% - 9%) were found exclusively within exons, despite the fact that exons have a double median size compared to introns (Spieth et al., 2014). A complete list of assigned peaks can be found in Table S2.

**Figure 3:**
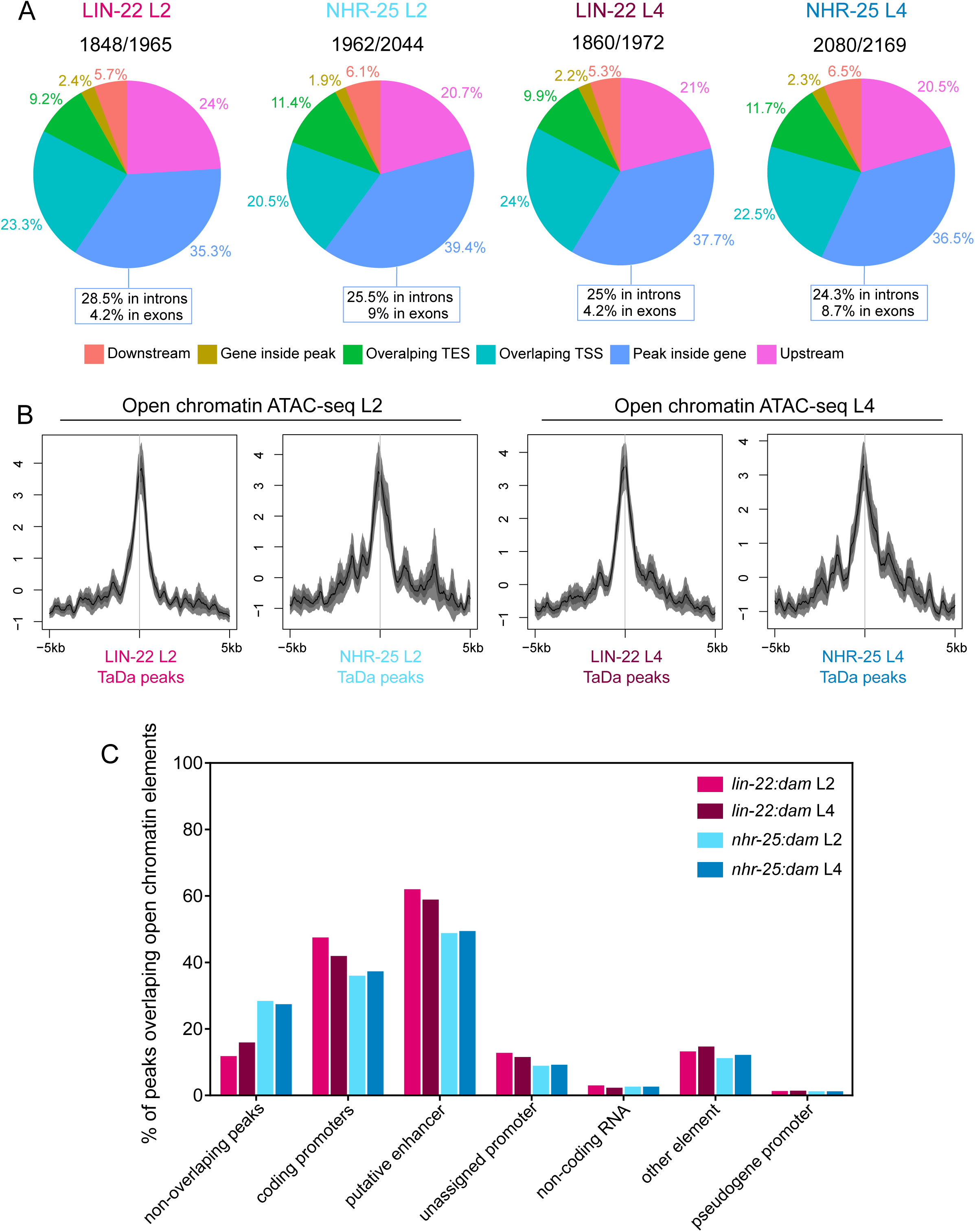
LIN-22 and NHR-25 peaks largely overlap genomic regions with regulatory potential. **(A)** Pie charts indicate the proportions of peaks residing in various genomic locations. Peaks were assigned to the single closest gene when their centre coordinate was found within 6 kb upstream and 1 kb downstream of the TSS and TES respectively of a gene. Ratios above pie charts show the number of assigned peaks to the total number of significant peaks found. The proportions of peaks within genes with exclusive intron or exon localisation are indicated under the pie charts. **(B)** Aggregation plots of open chromatin signal from ATAC-seq L2 and L4 data (Jänes et al., 2018) over LIN-22 and NHR-25 TaDa L2 and L4 peaks respectively. Note increased chromatin openness at the sites of peaks. Y-axes are z-scores for the plotted sequence length and shaded areas represent 95% confidence intervals. ±5 kb around the peak centres is shown. **(C)** Proportions of total peaks for each indicated TF at each stage overlapping with different categories of regulatory annotated open chromatin elements. A single TF peak may overlap with more than one open chromatin element.

To consolidate the link between the identified peaks and putative regulatory regions of the genome, we studied the overlap between our data and open chromatin signal tracks from ATAC-seq experiments performed on whole-animal L2 and L4 *C. elegans* (Jänes et al., 2018). TaDa peaks for both TF-Dam fusions showed increased average chromatin openness compared to neighbouring regions (Figure 3B). The genome-wide overlaps were highly significant (*p<* 0.0001 by Monte Carlo simulations) with only a small fraction (11% - 27.6%) of the total identified peaks for each TF and developmental stage not overlapping with an open chromatin element (Figure 3C). Instead, the majority of peaks were overlapping either coding promoters (35.2% - 46.7%) or putative enhancers (48% - 61.2%) (Figure 3C). We finally investigated the overlap between our data and tissue-specific open chromatin elements at the L4 stage described in a recent study (Serizay et al., 2020). We found significant overlap with hypodermally-enriched accessible chromatin (10.8% - 41.5%, *p<* 0.0001) in comparison to other tissues like neurons (3.4% - 2.4%) or the germline (0.8% - 1.7%), where the overlap was not significant. Taken together, the overall localisation profiles of TaDa peaks strongly support that TaDa signal is likely to reflect genuine LIN-22/NHR-25 binding sites.

### Comparisons of peak localisation profiles across methods and transcription factors

To compare our TaDa results against other methods, we used two published ChIP-seq datasets for NHR-25 binding at L1 and L2 stage from Shao *et al*., 2013 (L1) and Araya *et al*., 2014 (L2). Initial qualitative assessment of the signal tracks showed some promising agreement (Figure 4A). At a genome-wide level, aggregate signal of the *nhr- 25:dam* L2 and L4 over all ChIP-seq L1 and L2 peaks, or vice verca, exhibited strong overlapping localisation (Figure 4B), supporting the similarity of peak profiles across methods. Both the L2 and L4 *nhr-25:dam* peaks showed significant localisation overlaps by Monte Carlo simulations across the genome with the 683 peaks of the NHR-25 ChIP-seq L1 and the 5980 peaks of the ChIP-seq L2 datasets (Figure 4C). For example, we found a substantial overlap with the L2 ChIP-seq dataset, with approximately 37% (726 peaks) and 38.5% (835 peaks) of the total L2 and L4 *nhr- 25:dam* peaks overlapping respectively. It is of note that in 75% of the 726 *nhr-25:dam* L2 peaks overlapping the ChIP-seq L2 peaks, the overlap included the highest enriched GATC fragment of the TaDa peak.

**Figure 4:**
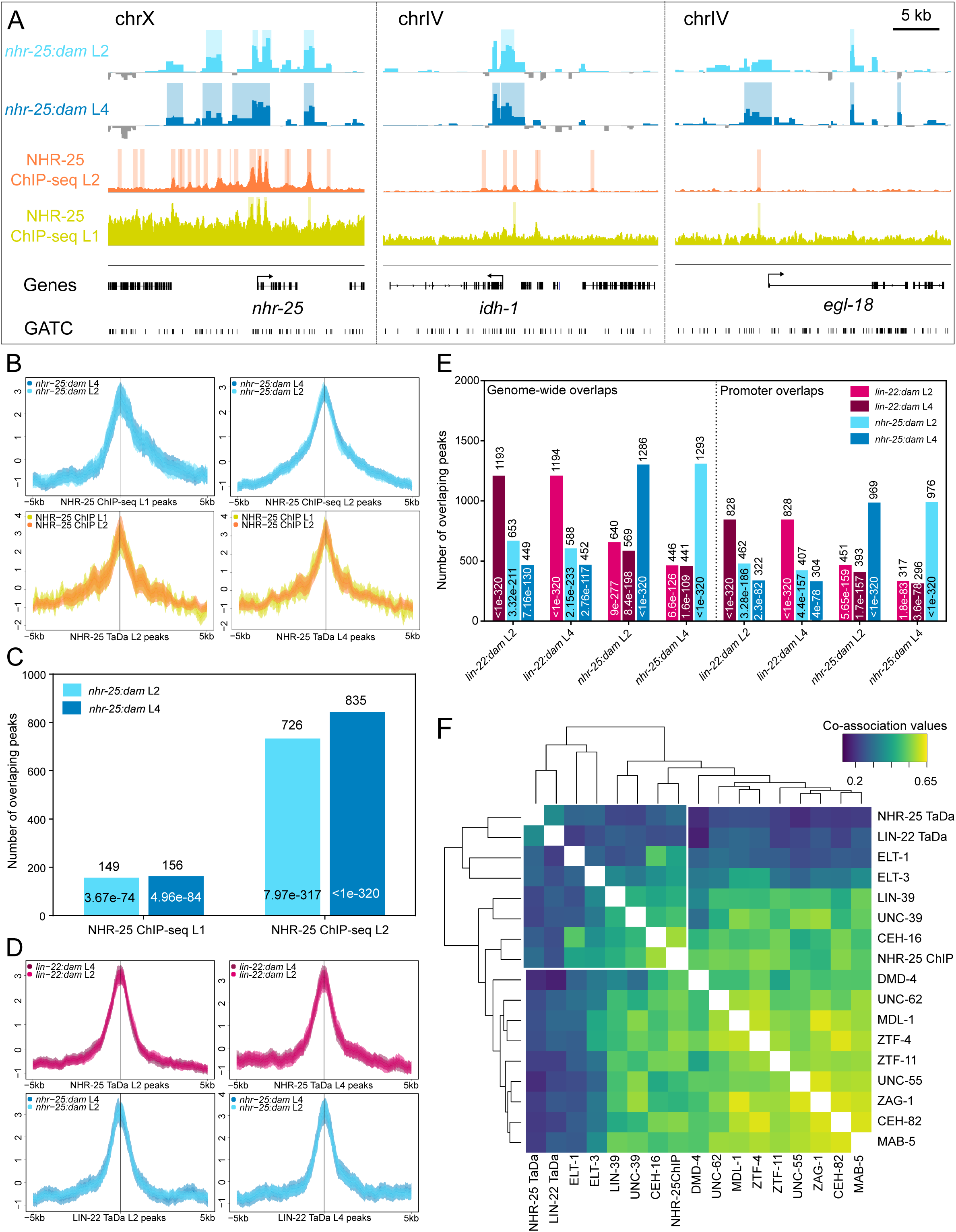
Peak localisation profiles align with ChIP-seq data and reveal overlap between the two transcription factors. **(A)** Representative snapshots showing agreement between *nhr-25:dam* TaDa data and two ChIP-seq signal profiles from L1 (Shao et al., 2013) and L2 (Araya et al., 2014) staged animals over *nhr-25*, *idh-1* and *egl-18*. Shaded regions indicate statistically significant peaks (FDR<0.05). The Y-axes for TaDa samples represent log_2_*(nhr-25:dam*/*NLS-GFP:dam)* scores with data range: -3 – 8 while for ChIP-seq the fragment pile-up per million reads score with data range: 0-10 for L1 and 0-1000 for L2. Scale bar is 5 kb as indicated. **(B)** Aggregation plots showing enrichment of *nhr-25:dam* L2 and L4 signal over centre of NHR-25 ChIP-seq L1 (left) and L2 (right) peaks and of NHR-25 ChIP-seq L1 and L2 signal over centre of NHR-25 TaDa L2 (left) and L4 (right) peaks. **(C)** Graph indicating the number of peaks from *nhr-25:dam* L2 and L4 samples that overlap with the L1 or with the L2 NHR-25 ChIP-seq peaks datasets. **(D)** Aggregation plots of *lin-22:dam* L2 and L4 signal over NHR-25 TaDa L2 and L4 peaks (top) and *nhr-25:dam* L2 and L4 signal over LIN-22 TaDa L2 and L4 peaks. All show increased signal for the peak location. **(E)** Graph showing the numbers of peaks for each colour-coded sample that overlap with the peaks of the sample indicated on the X-axis either across the whole genome (left) or restricted to promoter regions 5 kb upstream to 500 bp downstream of the TSS (right). **(F)** Hierarchical clustering of symmetrized co-association values for all pairwise comparisons between peak localisation patterns of the indicated TFs at the L2 stage. Peak profiles are taken from previous ChIP-seq experiments (Kudron et al., 2018) or our TaDa results. Note the separate clustering of TaDa peaks together with epidermal factors. In (B) and (D), ±5 kb around the peak centres have been plotted and the Y-axes represent z-scores for the plotted sequence length and shaded areas represent 95% confidence intervals. In (C) and (E), the exact number of peaks with overlaps is written above the bars and *p*-values from Monte Carlo simulations for the statistical significance of overlaps is shown inside the bar.

We then examined the overlap between the TaDa peak profiles identified for the two TFs. Interestingly, *lin-22:dam* signal showed an increased average preference to map onto NHR-25 TaDa peaks in comparison to adjacent regions and the same was true for *nhr-25:dam* signal on LIN-22 peaks (Figure 4D). Since LIN-22 and NHR-25 both participate in epidermal development, it is conceivable that this similarity may reflect genuine proximity or overlap of TF binding sites. Alternatively, the shared peaks could also reflect genomic sites where promiscuous binding of multiple TFs may occur. Such regions, usually referred to as High-Occupancy Target (HOT) regions have been previously determined in ChIP-seq experiments (Araya et al., 2014). TaDa peaks showed significant (*p*< 0.0001 by Monte Carlo simulations) yet smaller overlap with HOT regions compared to ChIP-seq (8% for *nhr-25:dam* as opposed to 34% forn NHR- 25 ChIP-seq L2 peaks). Furthermore, out of the 2167 HOT regions found in L2 (Araya et al., 2014), 13% were overlapping with TaDa NHR-25 L2 peaks, as opposed to 83% with NHR-25 ChIP-seq L2 peaks.

To further investigate the overlap between the TaDa profiles for the two TFs, all the pairwise comparisons were made and found to be non-random by Monte Carlo simulations. Around 60% of peaks of each profile overlapped across the two stages for the same TF, whereas <33% overlapped across TFs (Figure 4E left). To avoid *p*-value inflation when interrogating the entire genome while TF binding sites are expected to localise on promoters, overlaps and their statistical significance were re-calculated for promoter regions only (defined here as 5 kb upstream to 500 bp downstream of TSS). Again, we found promoter overlaps to be highly significant (Figure 4E right) and represented the majority of the genome-wide overlaps (67% - 72%). In addition, less than a third (24.3% -32.5%) of the overlaps in promoters across TF peaks occurred in HOT regions, suggesting that LIN-22 and NHR-25 may indeed share some targets in epidermal development.

To dissect the tissue-specificity of the binding profile of LIN-22 and NHR-25, we compared it to peak profiles for various TFs from ChIP-seq experiments conducted at L2 from the modERN database (Kudron et al., 2018). We included key TFs known to act specifically in the epidermis, such as ELT-1 and ELT-3, along with others that play broader roles in development including neurogenesis to investigate the global peak localisation pattern these TFs exhibit. Profile-wide comparisons of peak localisation between factors based on overlap and proximity statistics can provide a measure of co-association and statistically rank factors to highlight the broader similarity of their binding (Chikina and Troyanskaya, 2012). Previous calculations of co-association matrices have shown that TFs clustering together often regulate the same targets or act in the same tissue (Araya et al., 2014; Kudron et al., 2018). We found that our NHR-25 and LIN-22 TaDa datasets cluster together with the epidermal ELT-1 and ELT- 3 TFs, while ChIP-seq and TaDa profiles for NHR-25 did not cluster together (Figure 4F). Since NHR-25 also acts in non-epidermal cells, which is likely to be captured in ChIP-seq but not in the TaDa profiles, this may explain the distinct positioning of the NHR-25 binding profile depending on the method via which this profile was acquired (Figure 4F). These comparisons highlight the value of TaDa in revealing TF binding within a tissue of interest.

### LIN-22 and NHR-25 binding motif identification based on TaDa peaks

Next, we used the identified TaDa peaks to discover putative motifs that are associated to TF binding. Sequences restricted to the overlap between L2 and L4 peaks for each factor were used to increase the probability that these contain binding sites. The *de novo* identified motif for NHR-25 was consistent with the one previously reported by ChIP-seq and confirmed by functional studies (Araya et al., 2014; Barkoulas et al., 2016) (Figure S4A). When this motif was run against a database of known motifs (Fornes et al., 2020), it showed significant similarity to those of the *nhr-25* human orthologue NR5A1 (*p*=3.04×10^-6^) and mouse orthologue *Nr5a2* (*p*=2.35×10^-5^) (Figure S4B). The same analysis was carried out for LIN-22 and identified a motif that matches an E-box sequence (5’-CANNTG-3’) (Figure S4A), which matches reported motifs for the human orthologue HES1 (Lichtenberg et al., 2018). Comparison to known motifs showed significant similarity to that of the human HEY1 (Hes-related Family bHLH with YRPW Motif 1) factor (*p*=3.29×10^-3^), and HLH-1/ *MyoD* (*p*=2.45×10^-4^), which *Hes1* is known to antagonise for sites and binding partners (Sasai et al., 1992) (Figure S4B).

To assess if the identified motifs broadly represent preference for TF binding as determined in TaDa signal profiles, aggregate signal was mapped onto motif sites. For the NHR-25 motif, the *nhr-25:dam* L2 and L4 signal showed increased average preference for regions that include the motif as opposed to neighbouring sequences, while the *lin-22:dam* signal did not show any enrichment (Figure S4C). The reverse relationship was observed when the signal profiles were mapped onto the LIN-22 motif (Figure S4D). Some limited enrichment was observed for *nhr-25:dam* L2 data over the LIN-22 motif, which may be due to the fact that this motif is more noisy (Figure S4A), thus more frequent in the genome (1.5 times more frequent than the NHR-25 motif).

### Characterisation of LIN-22 and NHR-25 targets

To identify LIN-22 and NHR-25 putative targets, TaDa peaks were assigned to neighbouring genes. This assignment resulted in a set of 2809 genes for LIN-22 at L2 and 2833 genes at L4, and 3552 genes for NHR-25 at L2 and 3724 genes at L4 (Figure 5A, B and Table S2). The majority of the identified genes were shared between the two stages for each factor, with >63% of genes in any dataset being shared between L2 and L4 (Figure 5A, B). LIN-22 controls aspects of epidermal development by repressing neurogenesis and influencing ray formation in males (Wrischnik and Kenyon, 1997). These functions were reflected in related GO terms for the identified target gene sets at L2 and L4 (Figure 5C-E). In addition, the L2 dataset was found to be enriched for genes related to the Wnt signalling pathway (KEGG adjusted *p*-values of 3.28×10^-5^). The L2 and L4 datasets of putative NHR-25 targets were also enriched for multiple GO terms related to known NHR-25 biological functions (Chen et al., 2004; Chisholm and Hsiao, 2012; Gissendanner and Sluder, 2000; Hayes et al., 2006). For example, terms for structural constituents of the cuticle and molting cycle were found to be amongst the most significantly enriched (Figure 5F-H).

**Figure 5:**
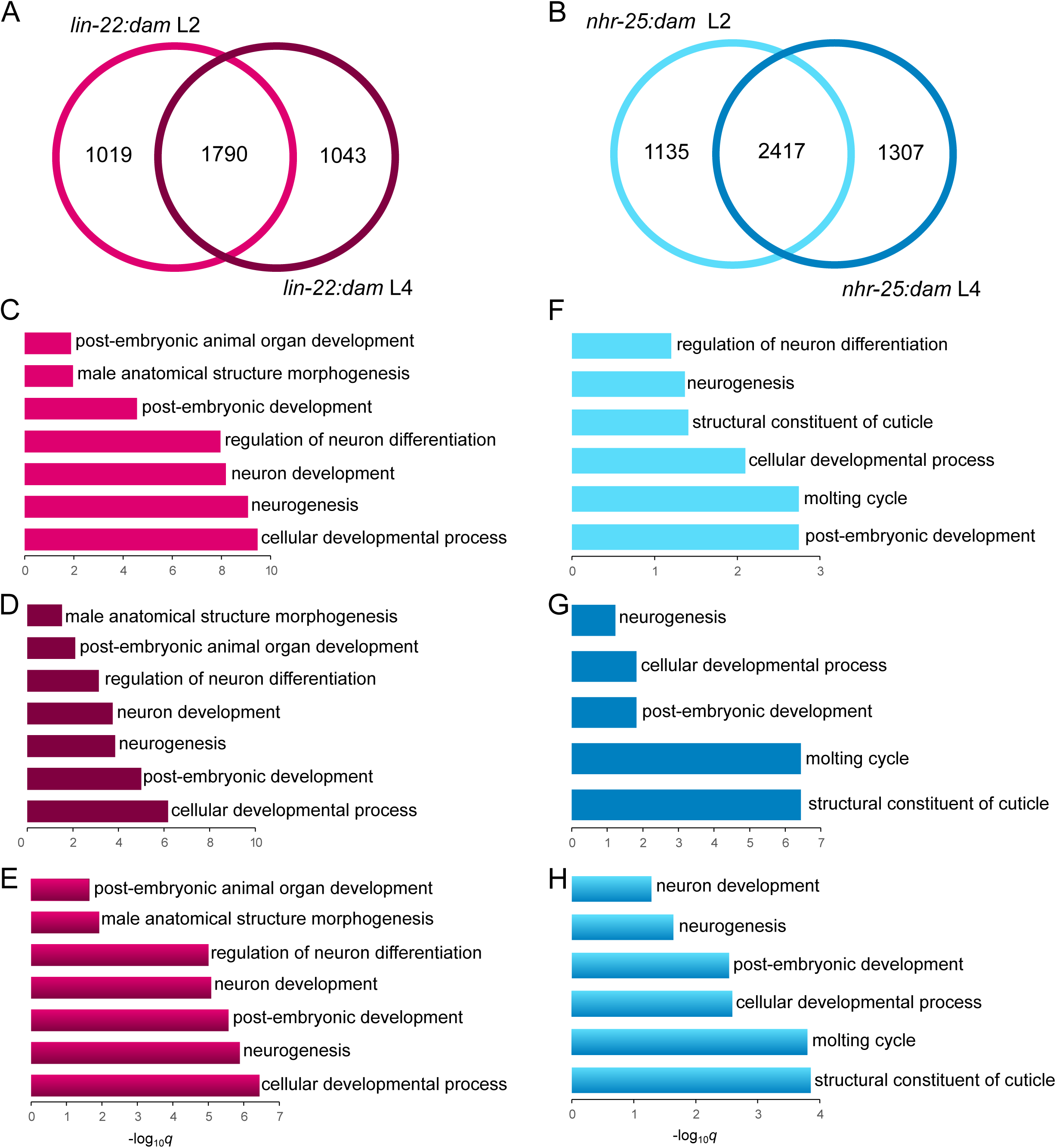
Enriched GO terms for LIN-22 and NHR-25 putative targets relate to their biological functions. **(A-B)** Venn diagrams for putative target intersections between genes identified at L2 and L4 for the *lin-22:dam* (A) and *nhr-25:dam* fusions (B). The intersection in both cases is significant with a hypergeometric distribution test *p* < 0.0001). **(C-H)** Plots of selected significantly enriched terms from GO term analysis for the genes in the L2 (C,F), L4 (E,H) or intersection datasets (D,G). For both TFs at all stages, relevant terms are recovered related to neurogenesis, development or molting.

We then sought to investigate how the genes identified via TaDa compare to those identified with other methods. To this end, we first compared the NHR-25 TaDa-identified targets with NHR-25 ChIP-seq datasets and found very significant overlap for all pairwise or higher order dataset intersections (Figure S5A). For example, the overlap with the ChIP-seq L2 dataset containing 7438 genes contained 62% of L2 and 61.6% of the L4 TaDa genes. We reasoned that some of the non-overlapping genes from ChIP-seq may reflect targets of NHR-25 outside the epidermis that are not captured by TaDa. Consistent with this hypothesis, tissue enrichment analysis for genes exclusively found by ChIP-seq showed higher enrichment for reproductive tissues compared to the epidermis (Figure S5B, C).

As there are currently no ChIP-seq datasets available for LIN-22, we intersected the TaDa-identified genes with *C. elegans* orthologs from ChIP-seq dataset for HES1, the human homologue of *lin-22*. We found statistically significant overlap containing genes that showed enrichment for the same GO terms previously identified in the TaDa gene sets alone, indicating conservation in regulatory interactions of LIN-22/HES1 across species (Figure S5D). A significant overlap was also found when the LIN-22 TaDa target genes were intersected with a list of 52 *in silico* predicted downstream genetic interactors of *lin-22* (Zhong and Sternberg, 2006). These included *lin-32* and *mab-5,* but also other genes known to participate in seam cell development like *unc-62* (Hughes et al., 2013), *nmy-2* (Ding and Woollard, 2017) or *rnt-1*, for which there was no prior evidence for regulation by LIN-22 (Figure S5D and Table S2).

We finally assessed the overlap between the identified targets for the two TFs. Interestingly, significant overlap was found in all pairwise comparisons across factors with 37%-46.4% of putative LIN-22 target genes overlapping NHR-25 datasets and 29.3%-35.3% of putative NHR-25 target genes overlapping LIN-22 datasets (Figure S5E). The intersection between all TaDa datasets contained 663 genes, which showed enrichment for GO terms related to neurogenesis and development (Figure S5E). These results indicate that the functions of the two TFs during epidermal development are likely to be executed by a combination of distinct and shared target genes.

### LIN-22 and NHR-25 targets suggest a link to cell differentiation

To validate putative interactions predicted by TaDa, we focused on selected candidates that are known to participate in the epidermal gene network. With regard to LIN-22, we focused on *rnt-1*, *cki-1* and *lin-17,* which showed significant signal enrichment in their promoter regions (Figure 2A and 6A). First, *rnt-1* is known to promote symmetric proliferative seam cell divisions (Nimmo et al., 2005). We found that *rnt-1* expression by smFISH is significantly increased in V1-V4 lineages in *lin-22(icb49)* null mutants, indicating that LIN-22 is likely to act as a repressor of *rnt-1* (Figure 6B, C). Second, *cki-1* encodes a cyclin-dependent kinase inhibitor protein (Buck et al., 2009; Hong et al., 1998). Comparison of *cki-1* expression between wild-type and *lin-22(icb49)* mutants revealed a significant reduction in transcript levels in V1-V4 lineages (Figure 6D, E). This finding suggests that LIN-22 may activate *cki-1* expression to modulate the differentiation programme.

**Figure 6:**
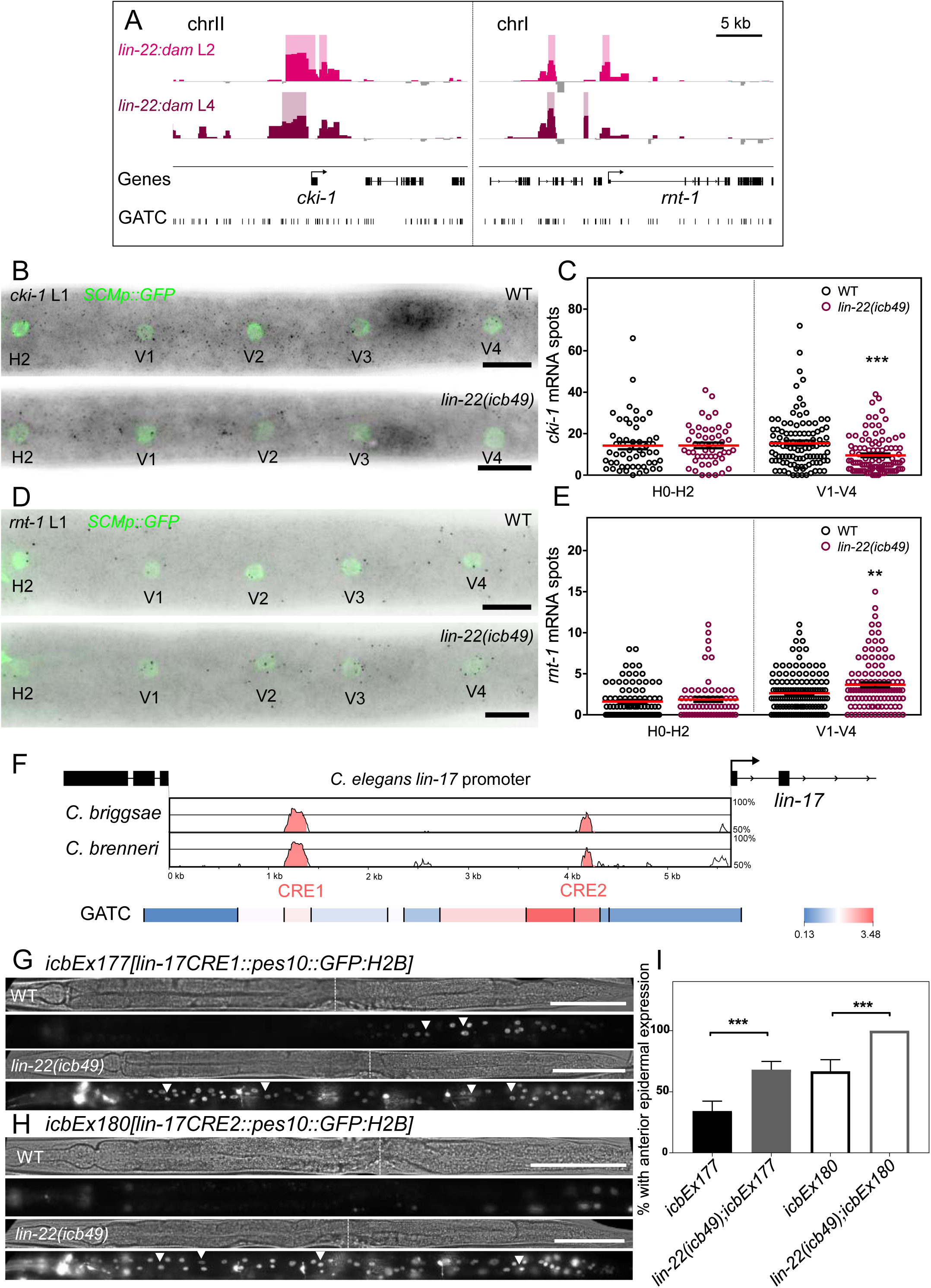
LIN-22 activates *cki-1* and represses *rnt-1 and lin-17.* **(A)** *lin-22:dam* signal forming significant peaks (shaded regions) around genes identified as putative LIN-22 targets. Y-axes are log_2_*(lin-22:dam*/*NLS-GFP:dam)* scores with data range: -1 – 3.5. **(B)** Representative *cki-1* smFISH images from WT and *lin-22(icb49)* animals at the late L1 stage. **(C)** Quantification of *cki-1* mRNA spots in H0-V4 seam cells, showing a significant reduction of expression in V1-V4 cells (50≤n≤126). **(D)** Representative *rnt-1* smFISH images from WT and *lin-22(icb49)* animals at the late L1 stage. **(E)** Quantification of *rnt-1* mRNA spots in H0-V4 seam cells, showing a significant increase in expression in V1-V4 cells (65≤n≤167). **(F)** Vista analysis of the *lin-17* promoter identified two conserved elements (*CRE1* and *CRE2*) between *C. elegans, C. briggsae* and *C. brenneri*. The position of the elements is shown on the *C. elegans* promoter sequence as a reference. Note that these elements are both overlapping GATC fragments with high TaDa enrichment scores (shown as red in the heatmap). **(G-H)** Representative brightfield and fluorescence images of L4 transgenic animals carrying transcriptional reporters for the *lin-17 CRE1* (*icbEx177* transgene) (G) and *CRE2* sequences (*icbEx180* transgene) (H) fused to *GFP:H2B* in WT and the *lin-22(icb49)* mutant background. Note that expression is restricted to few posterior cells in WT and expands to the anterior epidermis in the mutants. White arrowheads indicate epidermal GFP expression. *lin-22(icb49)* mutants also carry a *dat-1p::GFP* reporter labelling dopaminergic neurons, which is distinguishable by the visible axons. **(I)** Quantification of the proportion of transgenic animals with epidermal expression anterior to the vulva in WT (n=35 for *CRE1* and n=24 for *CRE2*) and *lin-22(icb49)* mutants (n=47 for *CRE1* and n=20 for *CRE2*). Error bars indicate the standard error of the proportion. Black stars show statistically significant differences with a Fisher’s exact test, *** *p*<0.001. In B and D, seam cells are labelled with *SCMp:GFP* and black spots correspond to investigated mRNAs. Scale bars are 5 kb in A, 10 μm in B and D, 100 μm in G and H.

Third, TaDa revealed two major sites of LIN-22 signal enrichment on the *lin-17* promoter (Figure 2A), which largely overlapped with two conserved regions (termed CRE1 and CRE2) between *C. elegans* and related *Caenorhabditis* species as revealed by Vista analysis (Frazer et al., 2004) (Figure 6F). To test the significance of these *cis*-regulatory elements, we constructed transcriptional reporters containing the CRE1 or CRE2 sequence fused to a minimal core promoter driving the expression of histone bound GFP. Multi-copy arrays were created for each element and introduced into the *lin-22(icb49)* mutant background. Both reporters were sufficient to drive expression in the posterior epidermis in wild-type animals, while expression expanded to more anterior epidermis in the *lin-22(icb49)* mutant background (Figure 6G, H, I). Taken together, these data indicate that LIN-22 is likely to bind to these regulatory elements on the *lin-17* promoter to repress *lin-17* expression. It is of note that LIN-22 was found to possibly target other Wnt receptors (*mom-5),* as well as Wnt ligands (*cwn-2*), Wnt secretion factors *(mom-1, mig-14*) and components of the signal transduction machinery (*lit-1*, *bar-1*) (Figure S6), indicating that LIN-22 may regulate the Wnt pathway at various levels of the signalling cascade.

With regard to putative targets of NHR-25, we focused on *egl-18* and *elt-1,* which showed signal enrichment in their promoter region (Figure 7A). These genes encode GATA TFs that promote seam cell fate and had not been previously linked to regulation by NHR-25. In particular, *egl-18* is a target of the Wnt signalling pathway that is known to maintain seam cell fate (Gorrepati et al., 2013; Koh and Rothman, 2001). Expansion of the *egl-18* expression domain to the anterior seam cell daughters that normally adopt the hypodermal differentiation programme has been shown to correlate with ectopic maintenance of the seam cell fate (Gorrepati et al., 2013; Katsanos et al., 2017). Furthermore, *elt-1* is thought to be the master epidermal fate regulator in *C. elegans* and is known to regulate *nhr-25* in the embryo (Chisholm and Hsiao, 2012; Gilleard and Mcghee, 2001). To test whether *elt-1* and *egl-18* are likely to be targets of NHR-25, we carried out smFISH quantifications in control and *nhr-25* RNAi treated animals at the L3 division stage. The efficacy of the RNAi treatment was confirmed by scoring for an increase in terminal seam cell number (Figure 7B). In *nhr-25* RNAi treated animals, an overall increase in *egl-18* expression in V1-V4 daughters was observed (Figure 7C, D), indicating that NHR-25 is likely to be a repressor of *egl-18.* Similarly, *elt-1* expression was found to be increased in anterior daughters of the V1-V4 lineages in *nhr-25* RNAi treated animals (Figure 7E, F). These findings suggest that NHR-25 promotes the differentiation programme by directly repressing key seam cell fate-promoting factors.

**Figure 7:**
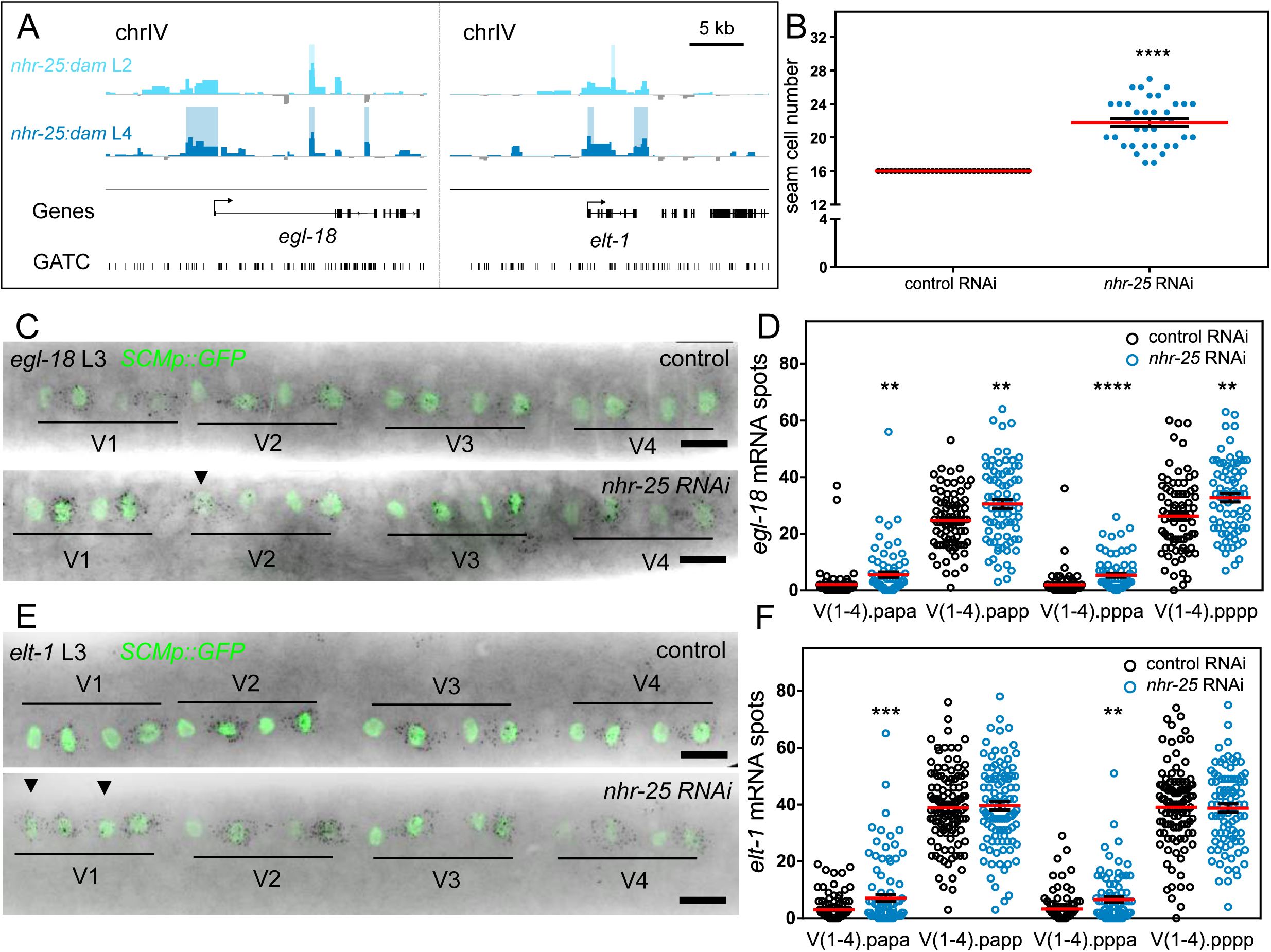
NHR-25 represses *egl-18* and *elt-1.* **(A)** *nhr-25:dam* signal forming significant peaks (shaded regions) around genes identified as putative NHR-25 targets. **(B)** Seam cell number comparison between control and *nhr-25* RNAi treated animals used for smFISH (n≥36). **(C)** Representative *egl-18* smFISH images of control and *nhr-25* RNAi treated animals during the L3 division. **(D)** Quantification of *egl-18* mRNA spots in the V1-V4 daughter cells following the L3 division (60≤n≤88) showing an overall increase in expression upon *nhr-25* RNAi. **(E)** Representative *elt-1* smFISH images of control and *nhr-25* RNAi treated animals during the L3 division. **(F)** Quantification of *elt-1* mRNA spots in the V1-V4 daughter cells following the L3 division showing a significant increase in expression in the anterior daughters (Vn.papa and Vn.pppa, 82≤n≤116). In C and E, seam cells are labelled with *SCMp:GFP* and black spots correspond to investigated mRNAs. Scale bars are 5 kb in A and 10 μm in C and E. Arrowheads in C and E indicate instances of strong expression in anterior daughter cells.

## Discussion

### LIN-22 and NHR-25 targets provide insights into epidermal cell fate specification

Our study refines the position of *lin-22* and *nhr-25* in the epidermal gene network. Based on our findings and previous literature (Brabin et al., 2011; Cassata et al., 2005; Chisholm and Hsiao, 2012; Hughes et al., 2013; Katsanos et al., 2017; Koh and Rothman, 2001; Thompson et al., 2016; van der Horst et al., 2019), we present an updated gene network underlying cell fate decisions in the epidermis (Figure 8). In light of our TaDa findings, we propose that LIN-22 and NHR-25 play a prominent role in mediating correct hypodermal differentiation in the epidermis. NHR-25 was known to influence seam cell patterning by regulating seam cell shape and establishment of cell-to-cell contacts post cell division (Chen et al., 2004; Gissendanner and Sluder, 2000; Šilhánková et al., 2005), although its link to cell fate specification was not well understood. Our findings suggest a role in the specification of the hypodermal cell fate through direct suppression of core seam cell specifying factors, such as *egl-18* and *elt-1*.

**Figure 8:**
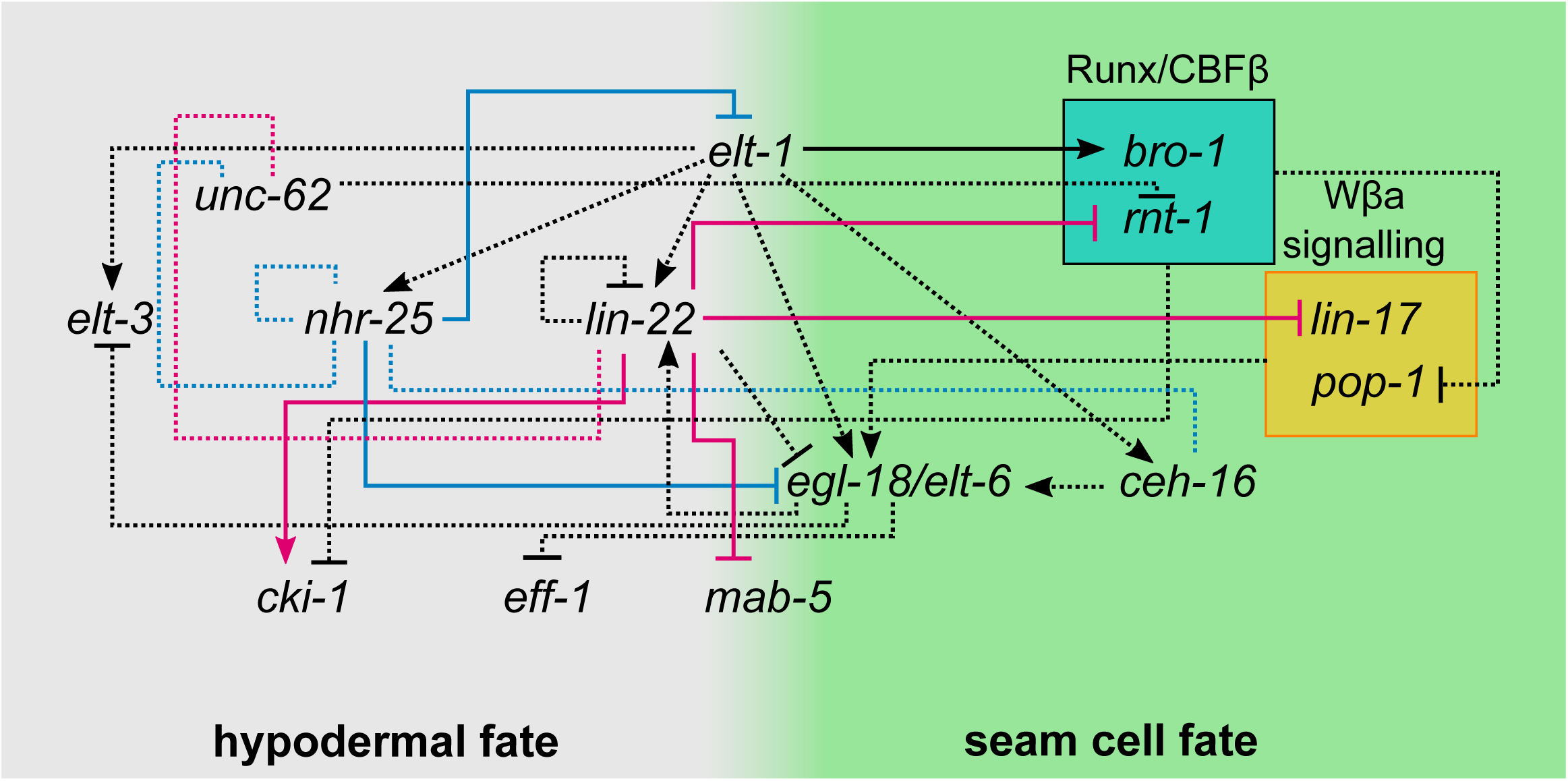
A combined gene network of epidermal cell fate interactions. Blue and magenta dashed lines indicate new interactions predicted by TaDa for LIN-22 and NHR-25 respectively. Activation or repression of targets is shown based on smFISH validation when known. Black dashed lines indicate published genetic interactions that is yet unknown whether they are direct or not (except for the link between *elt-1* and *bro-1*, which is likely to be direct). Hypodermal fate is shown in grey and seam cell fate in green.

We previously proposed that LIN-22 influences seam cell divisions by antagonising Wnt signalling through regulation of the expression of the Frizzled receptor *lin-17* (Katsanos et al., 2017). LIN-22 is also known to supress neurogenesis in V1-V4 seam cells through suppression of the Hox gene *mab-5,* as well as the pro-neuronal factor *lin-32* (Wrischnik and Kenyon, 1997). However, in all these cases, there was no prior evidence for a direct interaction. Our TaDa signal profiles reveal that LIN-22 is likely to directly bind to *mab-5* and *lin-32* upstream sequences in the seam cells. Although *Hes*-related factors are commonly associated with Notch signalling in other systems (Kageyama et al., 2007), we found an unusual link between LIN-22 and Wnt signalling in *C. elegans*, potentially through direct regulation of multiple components within this signalling pathway. LIN-22 was also found to repress *rnt-1,* the Runx homologue of *C. elegans,* which in complex with BRO-1 promote seam cell fate and symmetric divisions by repressing *pop-1* (van der Horst et al., 2019). This interaction is consistent with the observation of ectopic symmetric divisions in *lin-22* mutants (Katsanos et al., 2017). LIN-22 may activate the cell cycle inhibitor *cki-1*. This gene is expressed in the seam cells and *cki-1* RNAi increases seam cell number (Buck et al., 2009; Hong et al., 1998). Since *cki-1* drives G1 arrest allowing differentiation to progress (Hong et al., 1998; Matus et al., 2015), reduction of *cki-1* expression may contribute to the supernumerary seam cells observed in *lin-22* mutants. Canonical *Hes* factors are thought to generally act as repressors (Kageyama et al., 2007), for example the mammalian homologue of *cki-1*, p27^Kip1^ has been shown to be directly repressed by Hes1 in mice (Murata et al., 2005). In contrast to canonical HES factors, LIN-22 lacks a Groucho interacting domain (Schlager et al., 2006), so it may act as a repressor or activator depending on available co-factors or competition with other TFs for binding.

### Expanding the *C. elegans* toolkit for transcription factor target identification

We introduce in this study TaDa as a method to identify TF targets in a tissue-specific manner in *C. elegans*. Tissue-specific or constitutive levels of expression of Dam-fusions have been known to lead to toxicity, which we also confirmed here in the case of the *C. elegans* epidermis (Schuster et al., 2010; Southall et al., 2013; van Steensel and Henikoff, 2000). The TaDa transgene configuration overcomes this obstacle by minimising the levels of Dam expression in the tissue of interest. We show that TaDa can be used in early larval stages and dividing cell populations, such as the seam cells at the L2 stage, while still leading to robust identification of putative targets.

Currently, ChIP-seq is the most commonly used method to identify TF targets in *C. elegans*, with many datasets available based on large-scale projects (Celniker et al., 2009; Kudron et al., 2018). We present here evidence that TaDa is comparable to ChIP-seq in identifying target genes, while it offers two key advantages. First, TaDa allows the identification of TF targets in a specific tissue of interest without cell isolation, as opposed to all tissues where the TF is expressed (Aughey et al., 2019; Aughey and Southall, 2016). Second, TaDa requires substantially less material than ChIP-seq. We found approximately 2000 animals to be sufficient to allow the recovery of enough methylated DNA from the seam cells, which represent 3% of the total cells of *C. elegans*. This is also consistent with observations in mammalian cell lines, where a lot more cells were used to identify targets for the same TF by ChIP-seq than TaDa (Cheetham et al., 2018).

When direct comparison across methods could be made, the majority of TaDa putative target genes (>61%) were also identified by stage-matched ChIP-seq. The NHR-25 ChIP-seq L2 dataset was approximately three times larger in terms of peaks compared to TaDa L2 (2044 peaks in TaDa compared to 5980 in ChIP-seq). Although tissue specificity is likely to contribute to this difference, there are also alternative explanations. Both ChIP-seq and DamID methods have inherent biases due to the protocol and reagents used including the quality of the Dam fusions (Ramialison et al., 2017), issues with antibody affinity or PCR biases (Meyer and Liu, 2014). A key limitation in all DamID-based methods is the dependence on availability of GATC sites, which could hinder detection of some targets. However, the average length of GATC fragments in *C. elegans* is 367 bp and the median 204 bp, so depletion of GATC sites is unlikely to be pervasive enough to significantly undermine detection of targets. TaDa peak profiles may be distinct to ChIP-seq in other ways too, for example they were found to be less inclusive of HOT regions (Teytelman et al., 2013), with only 13% of L2 HOT regions being represented in NHR-25 TaDa in contrast to 83% in ChIP-seq. While TaDa peaks tend to be broader than those obtained from ChIP-seq experiments, we demonstrate that they can be used to identify TF binding motifs, for example based on GATC fragments that show the highest local enrichment.

In summary, TaDa can be a powerful approach to dissect complex TF behaviours related to tissue-specific target binding. This includes for example the HLH family of transcription factors in *C. elegans* that are thought to bind different targets in different tissues depending on their dimerising partners (Grove et al., 2009). TaDa predictions can be followed up with single cell resolution using smFISH in appropriate mutant or silenced backgrounds. This experimental combination can be useful to add quantitative strengths to newly described interactions, thereby facilitating mathematical modelling of developmental gene networks (Piano et al., 2006). The revised epidermal network represents a framework for future experiments to build upon. Given the conserved nature of some of the participating factors, this network can inform about interactions that underlie more broadly stem cell behaviour to dissect the developmental logic of robust stem cell fate patterning.

## Acknowledgments

We thank Tony Southall, Colin McClure and Gabriel Aughey for discussions on setting up the method. We thank members of the Barkoulas lab and David Matus for comments on the manuscript. Some *C. elegans* strains were provided by the CGC, which is funded by NIH Office of Research Infrastructure Programs (P40 OD010440). We acknowledge the support from the European Commission [ROBUSTNET-639485]. DK was the recipient of an Imperial College President’s PhD scholarship.

## Author contributions

DK carried out all experiments and data analysis. MB supervised the work. DK and MB wrote the manuscript.

## Materials and methods

### *C. elegans* maintenance

The *C. elegans* strains used in this study were maintained on standard Nematode Growth Medium (NGM) plates and were grown monoxenically on a lawn of *E. coli* OP50 at 20 °C. For TaDa experiments, strains were grown for at least two generations on a lawn of a *dam^-^/dcm^-^ E. coli* mutant of the K12 strain (New England biolabs, C2925). The laboratory reference N2 strain was used as the reference strain in this study. A complete list of strains used in this study is available in Table S3.

### Molecular cloning

To construct a TaDa backbone plasmid for epidermal expression of TFs fused upstream of Dam, the pCFJ151 universal MosSCI vector (Frokjaer-Jensen et al., 2014) was digested with BcuI/BspTI enzymes. The promoter of *wrt-2* was amplified from N2 lysate with oligos PB16 and PB7, the *C. elegans* optimised *mCherry* was amplified from the pAA64 (Barkoulas et al., 2016) plasmid using oligos PB8 and PB17, the *Dam* sequence was amplified from the pUAST attB LT3 Dam plasmid (gift by Tony Southall) using oligos PB18 and PB13 and the *unc-54 3’ UTR* was amplified from N2 lysate using oligos PB14 and PB15. All 4 fragments and the digested backbone were inserted in a multi-fragment Gibson assembly reaction to produce the pPB7(*wrt-2p::mCherry::Dam::unc-54 3’UTR + cb-unc-119*) plasmid. The XmaJI site between *mCherry* and *Dam* was digested to linearise the vector and allow the in-frame to *Dam* insertion of the *nhr-25* coding sequence amplified using oligos PB19 and PB20 from N2 cDNA to produce the pPB10(*wrt-2p::mCherry::nhr-25:Dam::unc-54 3’UTR + cb-unc-119*) plasmid. To construct the seam cell driven *lin-22:Dam* fusion and the *NLS-GFP:Dam* control for the TF TaDa experiments, the *lin-22* gene was amplified with oligos DK11 and DK12 from fosmid WRM0627dG07 while *NLS-GFP* was amplified from plasmid pPD93_65 (Fire Lab vector Kit) using oligos DK15 and DK16. Both amplicons were inserted upstream and in-frame with *Dam* by Gibson assembly in an XmaJI linearised pPB7 vector like above. The resulting plasmids produced were pDK4(*wrt-2p::mCherry::lin-22:Dam::unc-54 3’UTR + cb-unc-119*) and pDK8(*wrt-2p::mCherry::nhr-25:Dam::unc-54 3’UTR + cb-unc-119*). To test the importance of the *mCherry* primary ORF to the viability of animals and methylation levels, versions of the *lin-22:Dam* and *NLS-GFP:Dam* TaDa constructs without *mCherry* were produced. In more detail, the pPB7 plasmid was digested with BcuI/MunI, the 4085 bp and 6141 bp fragments were excised, extracted and kept. *lin-22* was amplified from pDK4 using DK102 and DK11. The 2 digestion fragments, *the lin-22* amplicon and the repair oligo DK103 were all inserted into a Gibson reaction to produce pDK49(*wrt-2p::lin-22:Dam::unc-54 3’UTR + cb-unc-119*) which was then digested with BcuI/XmaJI to remove *lin-22* and insert via Gibson assembly a DK108 and DK15 amplified fragment of NLS-GFP from pDK8 to generate pDK50(*wrt-2p::NLS-GFP:dam::unc-54 3’UTR + cb-unc-119*).

For ease of future applications, a versatile TaDa vector called pDK7 was constructed. Briefly, the *att* recombination cassette of pDest R4-R3 (Invitrogen) was amplified including the *attR4* site, *ccdb* and *CamR* genes but excluding the *attR3* site using the oligos DK17 and DK18 including half of the *attL1* site sequence on the 3’ DK18 primer. *mCherry* was amplified from pPB7 using oligos DK19 and DK20 carrying the other half of the *attL1* site on the 5’ of DK19. *dam* was amplified from pPB7 with oligos DK21 and DK22. The *unc-54 3’UTR* was amplified from pPB7 with oligos DK23 and DK24. All 4 fragments were inserted in a Gibson assembly reaction along with BcuI/BspTI doubly digested pCFJ151 vector to generate pDK7(*attR4-L1::mCherry::Dam-myc::unc-54 3’UTR + cb-unc-119*).

For the *lin-17 CRE1* and *CRE2* transcriptional reporters the oligos DK115 and DK116 along with DK118 and DK119 were used to amplify each of the respective regions from N2 lysate. The *Δpes-10* core promoter was amplified from L3135 using either the *CRE1* or the *CRE2* compatible forward primers DK117 and DK120 along with the DK107 reverse and was cloned along with the respective CRE amplicon in a NheI/XmaJI digested pDK16 to create pDK59(*lin-17CRE1:: Δpes-10::GFPo-H2B::unc-54 3’UTR + cb-unc-119*) and pDK60(*lin-17CRE2:: Δpes-10::GFPo-H2B::unc-54 3’UTR + cb-unc-119*). A complete list of the oligos used in this study is presented in Table S4.

### Single copy transgenesis

Single-copy transgenic lines were produced using the *Mos1-*mediated single-copy insertion (MosSCI) method (Frokjaer-Jensen et al., 2014). Briefly, ∼30 day-one adult animals of the EG6699 strain with a *Mos1* transposon insertion on chromosome II (*ttTi5605* locus) showing the uncoordinated (Unc) phenotype were injected for each transgene insertion. All the MosSCI injection mixes comprised of 50 ng/μl of a universal MosSCI vector carrying the transgene of interest flanked by the *ttTi5605* left and right recombination arms along with plasmids harbouring the *Mos1* transposase (pCFJ601 at 50 ng/μl), a heat-shock inducible *peel-1* toxin (pMA122 at 10 ng/μl) and co-injection markers (pGH8 at 5 ng/μl, *myo-2::dsRed* at 2.5 ng/μl, *myo-3::mCherry* at 5 ng/μl). Post-injection, animals were kept at 25 °C until plates were completely starved. The heat-shock treatment that follows was performed at 34 °C for 3.5 hours, after which the plates were allowed to recover for 3 hours at room temperature before “reverse chunking” was performed, where NGM chunks from the lawn of a new plate were placed on top of the starved, treated plate with the OP50 lawn facing upwards (O’Connell, 2010). The next day the top of lawns were screened for *unc-119(-)* rescued animals with absence of co-injection markers, which were transferred on different NGM plates per injected P0. Homozygous lines were confirmed molecularly for the presence of single-copy transgene insertions using oligos NM3880 and NM3884. A complete list of the transgenes produced for this study along with injection mix make-up information is available in Table S5.

### Microscopy

For seam cell imaging, live animals were mounted on fresh 2% agarose pads containing 100 μM NaN_3_ for immobilisation on glass slides. The slides were then imaged using either an AxioScope A1 (Zeiss) upright epifluorescence microscope with a Light Emitting Diode (LED) light source fitted with a RETIGA R6™ camera (Q IMAGING) controlled via the Ocular software (Q IMAGING) or on an inverted Ti-eclipse fully motorised epifluorescence microscope (Nikon) with a metal halide light source fitted with an iKon M DU-934, 1024 x 1024 CCD-17291 camera (Andor) controlled via the NIS-Elements software (Nikon). Scoring of the terminal seam cell or postdeirid neuron number phenotype was performed in late-L4 or early adult animals, carrying the *SCMp::GFP* or *dat-1p::GFP* markers. The lateral side most proximal to the objective was counted for every animal.

To perform smFISH, large populations of animals were synchronised by bleaching and were subsequently grown at 20 °C for 18 hours to reach the late L1 and 35 hours for the L3 asymmetric division stage. Animals were fixed in 4% formaldehyde (Sigma-Aldrich) in 1x PBS (Ambion) for 45 min on a vertical Stuart™ Rotating disk (Cole-Palmer). They were washed with 1.5 ml 1x PBS twice before being stored at 4 °C in 70% Ethanol for at least 24 hours for permeabilisation. Hybridisation was performed in 100 μl of buffer (100 mg/ml dextran sulphate (Sigma-Aldrich), 10% formamide in 2x SSC) at 30 °C for 16 hours with1 μl of a diluted in water probe added. The probe dilution was between 1:5 and 1:50 of a custom-made mixture of 21-48 Cy5-labelled oligonucleotides (Biomers) targeting the gene of interest. A complete list of probes and their dilution used in this study along with their sequences is available in Table S4. Animals were washed in a solution of 10% formamide, 2x SSC, stained with DAPI and resuspended for imaging in 100 μl GLOX buffer (0.4% glucose, 10 mMTris-HCl in 2x SSC) supplemented with 1 μl of 3.7 mg/ml glucose oxidase (Sigma-Aldrich) and 1 μl of 5 mg/ml Catalase (Sigma-Aldrich). Imaging was performed using the Nikon set-up described above using the seam cells closest to the objective lens as homing coordinates to acquire 17 Z-stack slices with a step of 0.8 μm for each of the DAPI, Cy5 and GFP channels (Semrock). Acquisition was performed using a 100x oil immersion objective with exposure set at 100 ms for DAPI at 1/32 reduced light intensity, 3 s for Cy5 and 300 ms for GFP. Analysis was performed using a custom MATLAB (MathWorks) pipeline (Barkoulas et al., 2013). In brief, selected animal DAPI and GFP images were used to annotate seam cells and draw regions of interest (ROIs) around the nuclei for at least 5 slices within which smFISH spots would be counted. An animal specific threshold for spot detection was set by manually sampling spots.

### TaDa wet lab protocol

Strains for TaDa experiments were transferred onto *dam^-^/dcm^-^* plates by spot bleaching and two biological replicates in separate plates were processed simultaneously. For each strain and replicate, nine 55 mm plates fully populated by gravid adults were used for large-scale egg preparation with isolated embryos seeded in new *dam^-^/dcm^-^* plates that were incubated at 20 °C. Half of the resulting synchronised populations were collected in a 15 ml centrifuge tube after 24 hours at the L2 stage and the other half after 48 hours at the L4 stage using 2 ml M9 buffer per plate. All collected populations underwent extensive washing by centrifuging animals at 1200 g for 3 min, removing the supernatant and washing the pellet (∼100 μl) with 10 ml M9 at least 5 times. Animal pellets were frozen at -20 °C before gDNA extraction. For gDNA extraction pellets were lysed using 750 μl of Cell Lysis Solution (QIAGEN) containing 100 µg/ml of Proteinase K on a heat-block at 55 °C shaking at 500 RPM for 16 hours overnight. Lysates were treated with 4 μl of 100 mg/ml RNase A (QIAGEN) at 37 °C shaking at 500 RPM for 3 hours. In turn, 250 μl of Protein Precipitation Solution (QIAGEN) was added to each sample and were incubated on ice for 5 min followed by vigorous vortexing for at least 30 sec, another 5 min incubation on ice and a centrifugation at 6000 g for 10 min at 4 °C. The supernatant for each sample was treated with isopropanol, washed with 70% ethanol and the pellet was air-dried for 1 hour. DNA pellets were hydrated with 55 μl of UltraPure™ distilled water and were left for 48 hours at 4 °C for the DNA pellet to dissolve.

For isolation and amplification of the GATC methylated gDNA fragments the protocol followed here is an adapted version of the one presented in (Marshall et al., 2016) for TaDa in *Drosophil*a with minor alterations. Of the extracted gDNA samples above, a total amount of 5 μg was transferred into a new 1.5 ml tube and was brought to 43 μl with the addition of UltraPure™ Distilled Water. For those samples where 5 μg were not available, 43 μl of the original sample were transferred. To each of those tubes 5 μl of 10x CutSmart Buffer (New England Biolabs) and 2 μl of DpnI restriction enzyme (New England Biolabs) were added and mixed by gentle flicking to prevent shearing of gDNA. The samples were incubated at 37 °C for 16 hours overnight and were in turn cleaned-up using the QIAquick PCR Purification kit (QIAGEN) and eluted with 40 μl of 50 °C water. 20 μl of each clean digestion product were split equally in two 0.2 ml PCR tubes (15 μl in each) and 4 μl of *Adaptor ligation buffer* (5x T4 DNA Ligase Buffer (New England Biolabs) and 10 μM of the dsAdR adaptor along with 1 μl (400 U) T4 DNA Ligase (New England Biolabs) were added in each. The samples were then incubated at 16 °C for 2 hours, followed by 10 min at 65 °C in a thermocycler. The double-stranded adaptor dsAdR was initially prepared by mixing equal volumes of 100 μM of the single stranded oligos AdRT and AdRb in a 1.5ml tube, immersing in a boiling-hot water-bath and letting it to cool down to room temperature to allow for gradual annealing. Following the adaptor ligation each sample was mixed with 20 μl of a 2x DpnII Digestion Buffer and 10 units (1 μl) of DpnII restriction enzyme (New England Biolabs) mastermix and were incubated for 3 hours at 37 °C. At this stage for each original sample two 40 μl digestion reaction products were available. For methylated DNA amplification by PCR each one of these products was mixed with 118 μl of *DamID PCR buffer* and 2 μl (10 units) of MyTaq™ DNA polymerase (Bioline) and were aliquoted at 40 μl in 4 different 0.2 ml PCR tubes. The *DamID PCR buffer* consisted of 1.36x MyTaq™ Buffer (Bioline) and 1.06 μM of the DamID PCR primer (Adr_PCR) that anneals on the adaptor sequence. In total for each original gDNA sample 8 PCR reactions were performed using the following cycling program:

Single cycle of steps 1-4: 72 °C for 10 min, 94 °C for 30 sec, °C for 5 min, 72 °C for 15 min

3 Cycles of steps 5-7: 94 °C for 30 sec, 65 °C for 1 min, 72 °C for 10 min

21 Cycles of steps 8-10: 94 °C for 30 sec, 65 °C for 1 min, 72 °C for 2 min

Final extension step: 72°C 5 min, Slow cool down to room temperature and storing at 10 °C.

<u>T</u>he number of cycles for steps 8-10 was increased from 17 described in (Marshall et al., 2016) to 21, which is more commonly used in previous DamID experiments in *C. elegans* (Askjaer et al., 2014). The 8 reactions per sample were pooled and cleaned-up using QIAquick PCR Purification kit (QIAGEN). To remove the adaptor sequences from the resulting PCR products up to 2.5 μg of product was transferred to a new 1.5 ml tube and was digested with AlwI restriction enzyme (New England Biolabs) at 37 °C for 16 hours. The products were cleaned-up again using the QIAquick PCR Purification kit. These final purified amplicons were sent to GENEWIZ for library preparation and Next Generation Sequencing (NGS) using the Illumina HiSeq platform.

### Calculation of TaDa signal profiles for LIN-22 and NHR-25 binding

FASTQ files representing single-end reads for each sample and replicate, were initially assessed using the fastq-stats perl script (available at https://github.com/owenjm/damid_misc/blob/master/fastq-stats) for uncut adaptors, primer dimer and internal GATC content as a post-sequencing quality control step for the wet lab executed protocol. The sequencing reads were mapped on the *C. elegans* genome, sequence alignment read-count maps were generated and normalised log_2_(*TF:dam/NLS-GFP:dam*) ratio scores were calculated per GATC fragment of the genome using the perl script damidseq_pipeline v1.4.5 (Marshall and Brand, 2015) (available at https://github.com/owenjm/damidseq_pipeline). The pipeline was used calling Bowtie 2 v2.3.4 (Langmead and Salzberg, 2012) for alignment to *C. elegans* bowtie indices from genome assembly WBcel235 (available from illumina iGenomes page), Samtools v1.9 (Li et al., 2009) for alignment manipulations and a GATC fragment file with the coordinates of all GATC fragments across the *C. elegans* genome in gff format, built from a WBcel235 FASTA file (available at https://www.ensembl.org/Caenorhabditis_elegans/Info/Index) using the gatc.track.marker.pl script (available at https://github.com/owenjm/damidseq_pipeline). The number of usable reads that have been acquired by the experiments presented here and map only once to the genome varied from over 6 million up to ∼30 million reads per sample with the genome coverage being between ∼9 and 45 times. From the 2 replicates per TF-Dam fusion and control-Dam fusion all pairwise genome-wide log_2_(*TF:dam/NLS-GFP:dam*) calculations were performed and averaged into a single signal profile of log_2_(*TF:dam/NLS-GFP:dam*) enrichment scores per GATC of the genome for each TF at each developmental stage. Genomic coordinate files produced and used throughout this study were converted between formats (BED, GFF, Bedgraph, BigWIg, Wig, GTF) using Excel, the <u>Convert betweenGTrack/BED/WIG/bedGraph/GFF/FASTA files</u> tool of the Galaxy powered GSuite Hyperbrowser (elixir) (at https://hyperbrowser.uio.no/hb/!mode=advanced) and the UCSC browser binaries bedGraphToBigWig, BigWigToBedGraph, bigWigToWig. BED and GFF signal and feature track files were visualised using the SignalMap NimbleGen software (Roche).

### Peak-calling and gene-assignment

Identification of statistically significant enriched peaks across the genome for each TF and developmental stage was performed using the perl script find_peaks (available at https://github.com/owenjm/find_peaks) with an FDR<0.05 and default settings with the averaged log_2_(*TF:dam/NLS-GFP:dam*) signal profiles as input. The output is a list of genomic interval coordinates for statistically significant peaks with a peak enrichment score and an FDR value. Significant peaks were assigned to genes using UROPA (Kondili et al., 2017) as a web tool (available at http://loosolab.mpi-bn.mpg.de/UROPA_GUI/) with Caenorhabditis_elegans.WBcel235.99.gtf (from http://www.ensembl.org/Caenorhabditis_elegans/Info/Index) as the genome annotation file. Peaks were assigned to genes on any strand when their centre coordinate was positioned up to 6 kb upstream of a gene start site or 1 kb downstream of the end site. The location of the peak relative to the gene was assigned based on the full length of the peak and the strand of the gene. To avoid discarding valid regulatory relationships and due to the compactness of the *C. elegans* genome no prioritisation was set for peaks when multiple assignments were available and all are reported.

For the NHR-25 L1 ChIP-seq dataset (Shao et al., 2013) used in this study for peak localisation comparisons, raw sequencing data (GEO number: GSE44710) were processed into significant peak profiles using MACS2 on Galaxy (https://usegalaxy.eu/). NHR-25 and input bedgraph signal tracks were inserted into the MACS2 bdgcmp tool (default: Poisson *p*value algorithm) to deduct noise and identify NHR-25 specific signal. The output was then used with the MACS2 bdgpeakcall tool (cutoff 1.0, min-length 200 and max-gap 30) to generate genome-wide profiles of significant peaks. Profiles representing different replicates were merged using bedtools merge with averaged heights for peaks that overlap.

### Pearson’s correlation and principal component analysis

Correlation between samples and reproducibility of replicates was assessed using the deeptools3 (Ramírez et al., 2016) multiBamSummary (--binSize 300) and plotCorrelation (--corMethod pearson, --whatToPlot heatmap, --skipZeros, -- removeOutliers) tools on Galaxy (https://usegalaxy.eu/). Principal component analysis was performed using the deeptools3 plotPCA tool on the multiBamSummary-calculated read count density summary matrices.

### Aggregation plots and heatmaps of signal localisation

Signal localisation preference around given genomic features presented as aggregation plots or heatmaps were generated using the SeqPlots GUI application (Stempor and Ahringer, 2016) with specific settings mentioned individually for each presented result. Aggregation plots represent signal averages for 10 bp bins in regions of varying but specified length around positional features of the genome. For genes all of their start and end coordinates, based on the largest transcript and used here as the transcriptional start site (TSS) and transcriptional end site (TES) of genes, are anchored to two positions of the X-axis and their genic sequence is pushed or stretched to a pseudo-length of usually 2 kb. For other features the midpoint coordinate is used to align all to the same position on the X-axis which then extents upstream and/or downstream of that region. For each position around the feature an average is calculated across all the features to generate the aggregation plot line with a shaded area representing the 95% confidence interval. When z-scores are presented on the Y-axes, those have been calculated as deviations from the mean signal seen across the plotted region. In heatmaps each line represents each occurrence of a genomic feature indicated on the X-axis along with a surrounding region of a given length also indicated on the X-axis. Colourscales indicate the positional enrichment score calculated as averages per 10 bp bins.

### Assessment of overlaps between sets of genomic intervals or gene sets

To identify overlapping peaks between samples or other genomic interval or features the bedtools intersect tool was used with settings dependent on the prospected outcome of the processing. To test if sets of genomic coordinates representing various features show statistically significant overlaps across the genome, Monte Carlo simulations have been performed using the python pipeline OLOGRAM, part of the gtftk package (Ferré et al., 2019). *p*-values are calculated based on the occurrence of intersections between intervals and overall length of overlap (in bp) across the genome. For statistical assessment of the level of association between patterns of peak (TaDa or ChIP-seq peaks) localisation across the genome for different TFs, the IntervalStats tool (Chikina and Troyanskaya, 2012) as part of the coloc-stats webserver (https://hyperbrowser.uio.no/coloc-stats/) was used. In brief, for the TFs used in this study ChIP-seq optimal IDR-thresholded peak coordinate files from L2 animals were downloaded from the modERN (Kudron et al., 2018) and modENCODE (Celniker et al., 2009) databases and were combined along with the L2 TaDa TF samples into a Gsuite of genomic tracks on coloc-stats. Each peak file was then used as query against the GSuite of reference sequences to calculate the IntervalStats statistic for co-localisation for all pairwise comparisons of peak coordinates. The values in the resulting comparison matrix representing comparisons between the same two TFs with different directionality (query-reference) were averaged to symmetrise the matrix and calculate the final co-association values that were plotted as a heatmap using the R package heatmap3 and hierarchical clustering. To assess the statistical significance of overlaps between sets of genes, hypergeometric distribution or Fisher’s exact tests were performed either on http://nemates.org/MA/progs/overlap_stats.html or using the R software package *SuperExactTest* (Wang et al., 2015) respectively. For both tests when sets of coding genes are compared the size of the sampling pool was set to 20191, the number of annotated coding genes in the WBcel235.99 release. Representation of overlaps was either in the form of Venn diagrams generated using http://bioinformatics.psb.ugent.be/webtools/Venn/ or in the form of the output of the *SuperExactTest* package.

### TF motif identification by TaDa

Identification of motifs from transcription factor TaDa peaks was done here using HOMER (Heinz et al., 2010). The top 200 peaks with the highest averaged enrichment score were used for the analysis. Peak interval files were used as input for the findMotifsGenome.pl script using the ce11 genome assembly, masking of the sequences and the option to analyse the size of sequences provided by the interval file (options: ce11 -size given -mask). The logos presented here for motifs identified using homer were generated after converting the homer positional weight matrix into a transfac matrix using the RSAT (Nguyen et al., 2018) Metazoa convert matrix tool (http://rsat.sb-roscoff.fr/convert-matrix_form.cgi) and importing to Weblogo3 (Crooks et al., 2004) (http://weblogo.threeplusone.com/) for logo drawing. Identification of similar known motifs to the *de novo* identified motifs was performed using the Tomtom tool of MEME-suite (Gupta et al., 2007). The default parameters were used and the interrogated motif matrices were compared against the JASPAR core 2018 non-redundant database.

### Gene set enrichment analysis

Gene sets identified in this study were assessed for enriched gene ontology (GO) terms or association with tissue specific expression using the worbase.org Enrichment Analysis tool (Angeles-Albores et al., 2016; David Angeles-Albores, Raymond Y. N. Lee, n.d.) (https://wormbase.org/tools/enrichment/tea/tea.cgi), using a *q*-value threshold of 0.1. For significant GO terms presented here the –log_10_*q* value is plotted. Association of gene sets with biological pathways was evaluated using the gProfiler gOst tool (https://biit.cs.ut.ee/gprofiler/gost) and a g:SCS calculated significance threshold of 0.05.

### RNAi

Knockdown of *nhr-25* was performed by feeding animals on lawns of bacteria expressing dsRNA targeting *nhr-25* (Source Bioscience). Bacteria were grown overnight and then seeded directly onto NGM plates containing 1 μM IPTG, 25 μg/ml ampicillin and 6.25 μg/ml tetracycline. 5 L4 animals of the strain of interest were transferred on RNAi plates and were allowed to lay progeny that were phenotyped at the stage of interest. Control treatments were performed in parallel, with precisely the same experimental conditions, by feeding animals on lawns of the same strain of HT115 bacteria containing an empty vector.

### Statistical analysis

Statistical analysis for comparisons between datasets that is not covered in the above paragraphs was performed using GraphPad prism 7 (www.graphpad.com). To test differences in the mean between seam cell scoring or smFISH counting datasets an unpaired two-tailed t-test was performed when the comparison was between two datasets and a one-way Analysis of Variance (ANOVA) was performed when multiple datasets were compared. One-way ANOVA was followed by a Dunnet’s post hoc test when the mean of multiple datasets was compared to that of a control. The significance threshold used throughout is *p*<0.05.

## Data availability

All raw sequence files and processed signal files have been deposited in the National Center for Biotechnology Information Gene Expression Omnibus (accession number: GSE163243).

## Supplementary Tables

Table S1. Genome coordinates of significant TaDa peaks for NHR-25 and LIN-22 at the L2 and L4 stage.

Table S2. Assigned genes per TaDa peak for NHR-25 and LIN-22 at the L2 and L4 stage.

Table S3. Strains used in this study.

Table S4. Primers used in this study (general oligos and smFISH probes).

Table S5. List of transgenes and injection mixes used in this study.

## Supplemental Figure Legends

**Figure S1: Assembly and assessment of *in vivo* methylation by TaDa fusions.**

**(A)** Graphic illustration of the design of a versatile *C. elegans* universal TaDa cloning platform with its key features. From left to right the plasmid contains: universal MosSCI recombination sites (ttTi5605_R and L), an LR attR4-attL1 Gateway cloning site for promoter insertion, an *mCherry* primary ORF followed by 2 x STOP codons, indicated in red, a frameshift in yellow and a unique XmaJI restriction site followed by *dam*, unique restriction cites for PaeI, PacI, NheI and an in-frame STOP codon prior to an *unc-54 3’UTR*. **(B)** Example of amplification products from methylated gDNA extracted from strains carrying the *dam* fusions described in this study and a WT dam^-^ strain fed on dam^-^ *E. coli.* Note that all samples except for the *dam^-^* strain show a pronounced 2 kb to 200 bp smear. Wells for all samples were loaded with the same volume of reaction product. Also note the marked increase in methylation observed in the absence of *mCherry* as a primary ORF. Strains grown on *dam^+^* OP50 also show extensive amplification due to methylated bacterial DNA.

**Figure S2: Validation of expression and functionality of TaDa transgenes.**

**(A)** Confirmation of single-copy transgene expression in the *wrt-2* domain (seam cells) using *mCherry* expression (primary ORF). Animals were imaged at the L4 stage and white arrows mark weak expression in the seam, as expected for single copy transgenes. **(B)** Overexpression of the *lin-22:dam* construct as a multi-copy transgene (*icbEx54*) in the *lin-22(icb38)* mutant partially rescues the supernumerary PDE neurons phenotype labelled by *dat-1p::GFP* (n≥31). **(C)** Overexpression of the *nhr-25:dam* fusion as a multi-copy transgene (*icbEx184*) in the *nhr-25(ku217)* mutant significantly increased the mean seam cell number (n≥31). Scale bars in A are 20 μm. In B and C error bars indicate standard error of the mean and black stars indicate statistically significant differences in the mean with a t-test (* *p*<0.05, **** *p*<0.0001).

**Figure S3: TaDa replicate reproducibility and fusion-dependent methylation.**

**(A-B)** Pearson correlation heatmaps based on normalised aligned read count maps for the LIN-22 (A) and NHR-25 (B) TaDa experiments. The correlation coefficient for each pairwise comparison is printed in each cell of the heatmaps. All Dam-fusions show strong replicate reproducibility and high within-fusion correlation, indicated by high correlation coefficients. Note that TF and control samples show low correlation between them and cluster separately. **(C)** Summary heatmap of Pearson correlations between all samples for the TF TaDa performed in this study. Note the high correlation between control samples from different experiments and low correlation with distinct separate clustering between the *lin-22:dam* and *nhr-25:dam* samples. Samples belonging to the LIN-22 or NHR-25 experiment are indicated with purple or blue font respectively. **(D)** Principal component analysis on normalised aligned read count maps for all samples shows distinct grouping between TF and control fusions. **(E)** Principal component analysis only on *lin-22:dam* and *nhr-25:dam* replicates shows distinct grouping between the two TF samples. Samples in D and E are colour-coded as per the shared key in the middle. **(F)** Example of averaged signal enrichment profiles for *lin-22:dam* and *nhr-25:dam* fusions across a 1 Mb region of chromosome I (locations of protein coding genes are indicated as black lines / boxes. Signal enrichment forms distinct peaks for each TF that show some similarity between stages. The Y-axes represent log_2_*(TF:dam*/*NLS-GFP:dam)* scores. Scale bar length is 2 Mb.

**Figure S4: NHR-25 and LIN-22 binding motifs identified using TaDa peaks.**

**(A)** *De novo* identified DNA motifs for NHR-25 (left) and LIN-22 (right) binding from TaDa peaks. These motifs were found in 41% of the LIN-22 L2, 33% of the LIN-22 L4, 47% of the NHR-25 L2 and 51% of the NHR-25 L4 total TaDa peaks, with the co-occurrence being statistically significant using Monte Carlo simulation (*p*-values < 0.0001 in all cases)**. (B)** Similar known motifs to the TaDa-identified NHR-25 (left) and LIN-22 (right) motifs are shown from available databases. **(C-D)** Aggregation plots of TaDa signal for all TFs and available stages over instances of the NHR-25 TaDa motif (C) and the LIN-22 TaDa motif (D). Both TFs show stronger preference for their respective motif compared to the alternative motif. Regions of ±5 kb around the peak centres have been plotted, the y-axes represent mean enrichment scores for the plotted sequences and shaded areas represent 95% confidence intervals.

**Figure S5: TaDa identified targets for NHR-25 and LIN-22 show significant overlaps between them and with other published datasets.**

**(A)** Barplot showing the size of all possible intersections between: *nhr-25:dam* identified target genes at L2/L4, NHR-25 ChIP-seq-identified target genes at L1 (Shao et al., 2013) and NHR-25 ChIP-seq peaks at L2 (Araya et al., 2014) assigned to genes using the same method used in this study. Selected enriched GO terms for gene sets from the large intersections between TaDa and ChIP-seq L2 are shown. **(B-C)** Tissue-enrichment analysis in the gene set of putative targets that are common between NHR-25 L2 ChIP-seq and TaDa (B) or exclusively identified by ChIP-seq (C). In both cases, the top 15 most overrepresented tissues are shown. **(D)** Barplot showing the size of all possible intersections between: the identified target genes for *lin-22:dam* L2 and L4, genes predicted to be downstream genetic interactors of *lin-22* (Zhong and Sternberg, 2006) and the *C. elegans* orthologues of HES1 targets based on ChIP-seq (Encode project accession: ENCSR109ODF). Selected enriched GO terms for gene sets from the intersections between orthologues of HES1 ChIP-seq targets and TaDa targets are shown. Genes in the pairwise intersection between TaDa and predicted interactors are listed in full above the bars. **(E)** Plot showing all possible intersections between TaDa-identified target genes for LIN-22 and NHR-25 at L2 and L4. Selected enriched GO terms are shown for the genes common in all datasets. In A, D the size of each individual gene set is printed at the bottom right of each graph. In A, D and E the statistical significance of each intersection assessed by a Fisher’s exact test is colour-coded as shown in the key (yellow, orange, red marks significant overlaps).

**Figure S6: TaDa reveals a link between LIN-22 and multiple components of the Wnt signaling pathway.**

Signal profiles from *lin-22:dam* showing enrichment forming statistically significant peaks (shaded regions) on sequences associating with the Wnt component-encoding genes *mom-1, mig-14, cwn-2, lit-1, bar-1* and *mom-5.* Y-axes are log_2_*(lin-22:dam*/*NLS-GFP:dam)* scores with data range: -1 – 3.5.

