## Supplementary figures and images for "Targeted DamID in *C. elegans* reveals a role for LIN-22 and NHR-25 in epidermal cell differentiation"

### Supl Figures 1-6

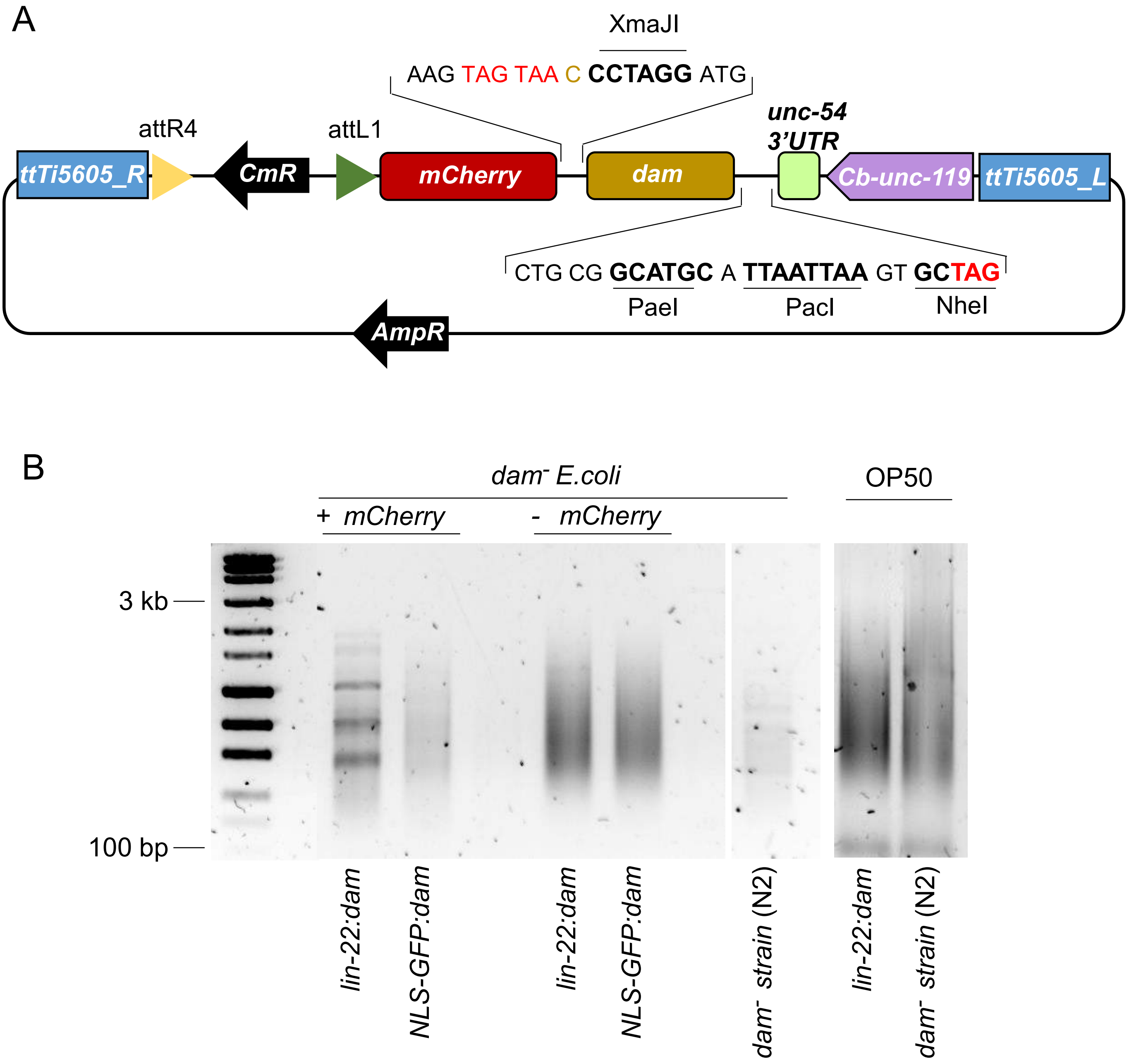

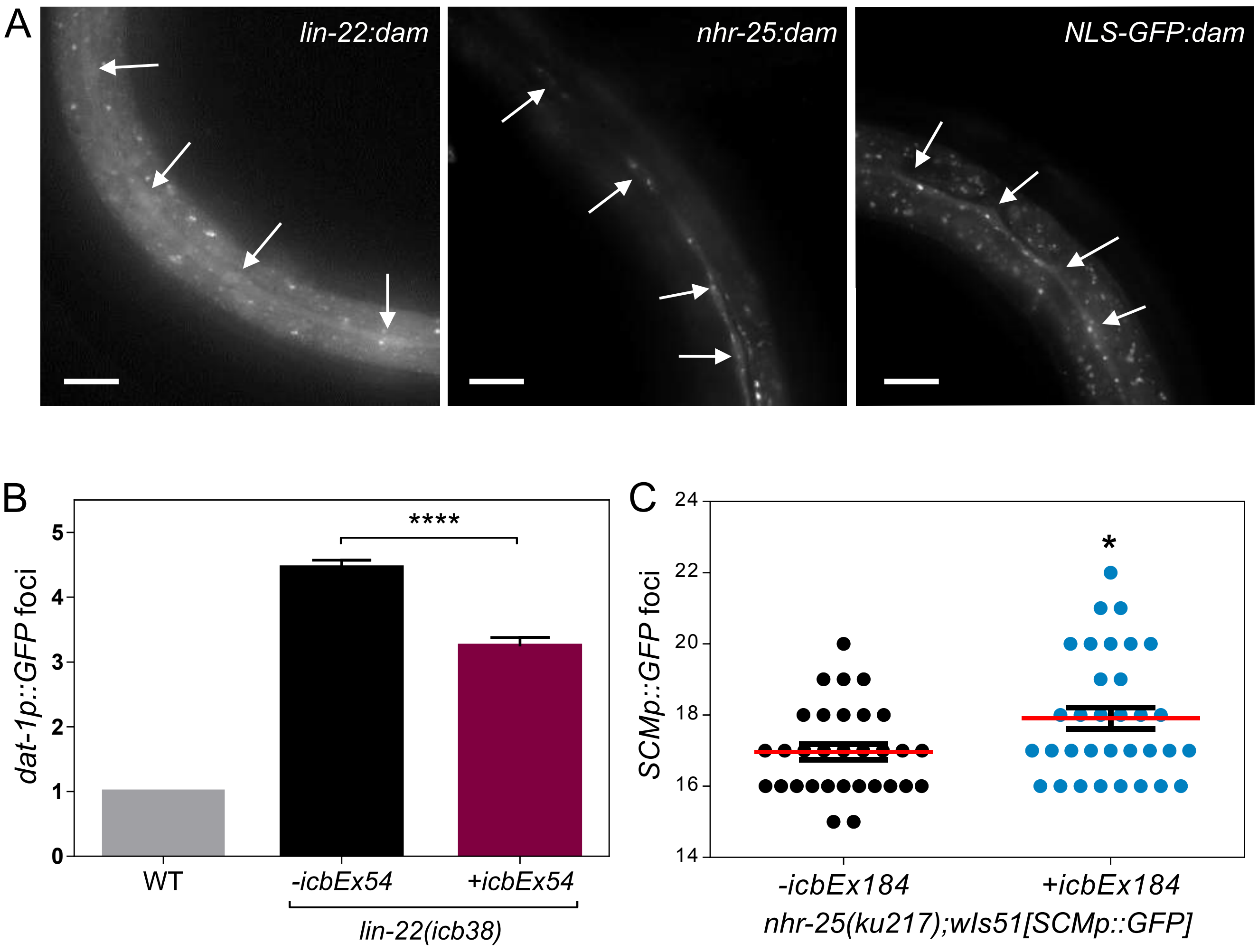

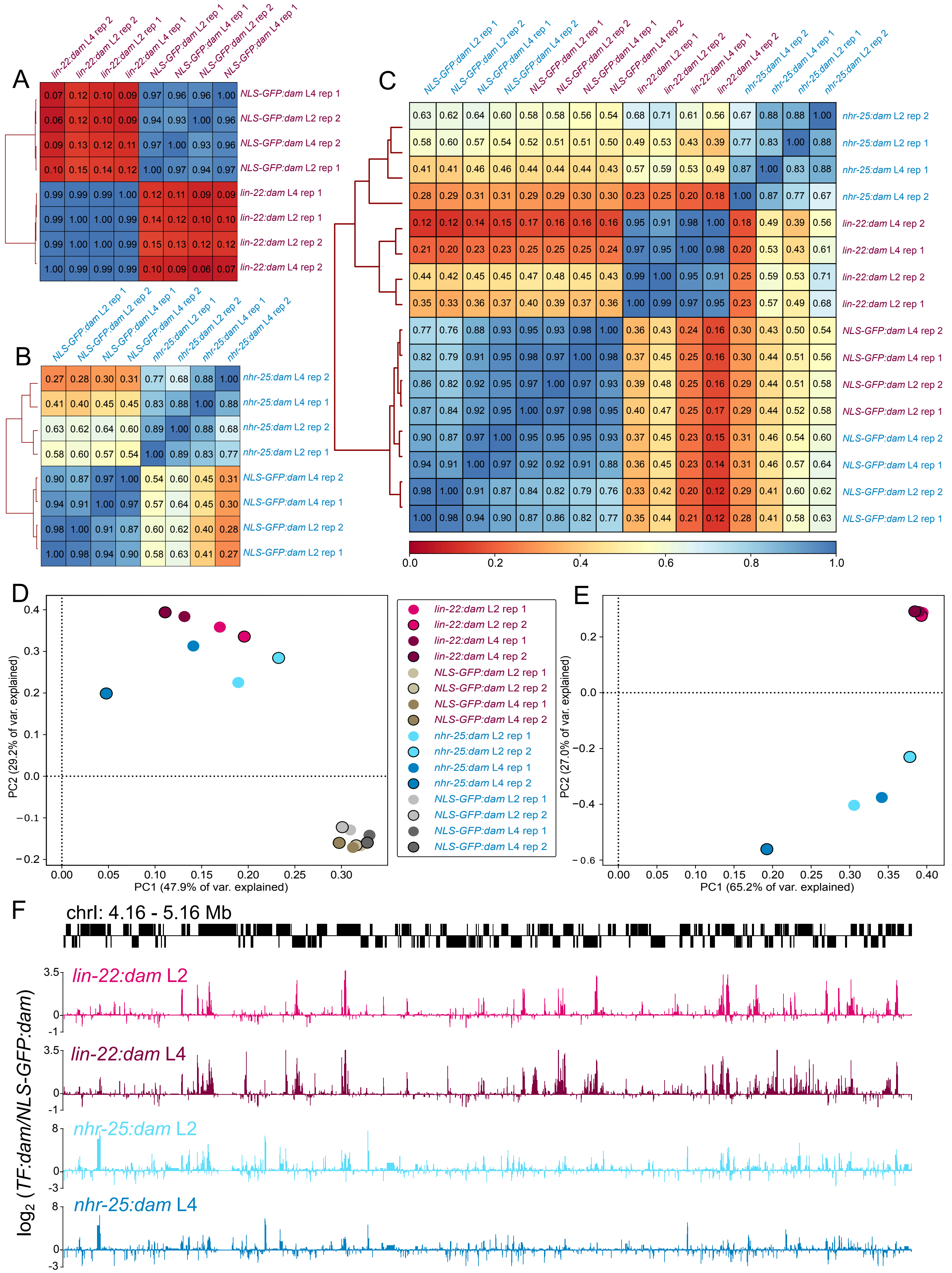

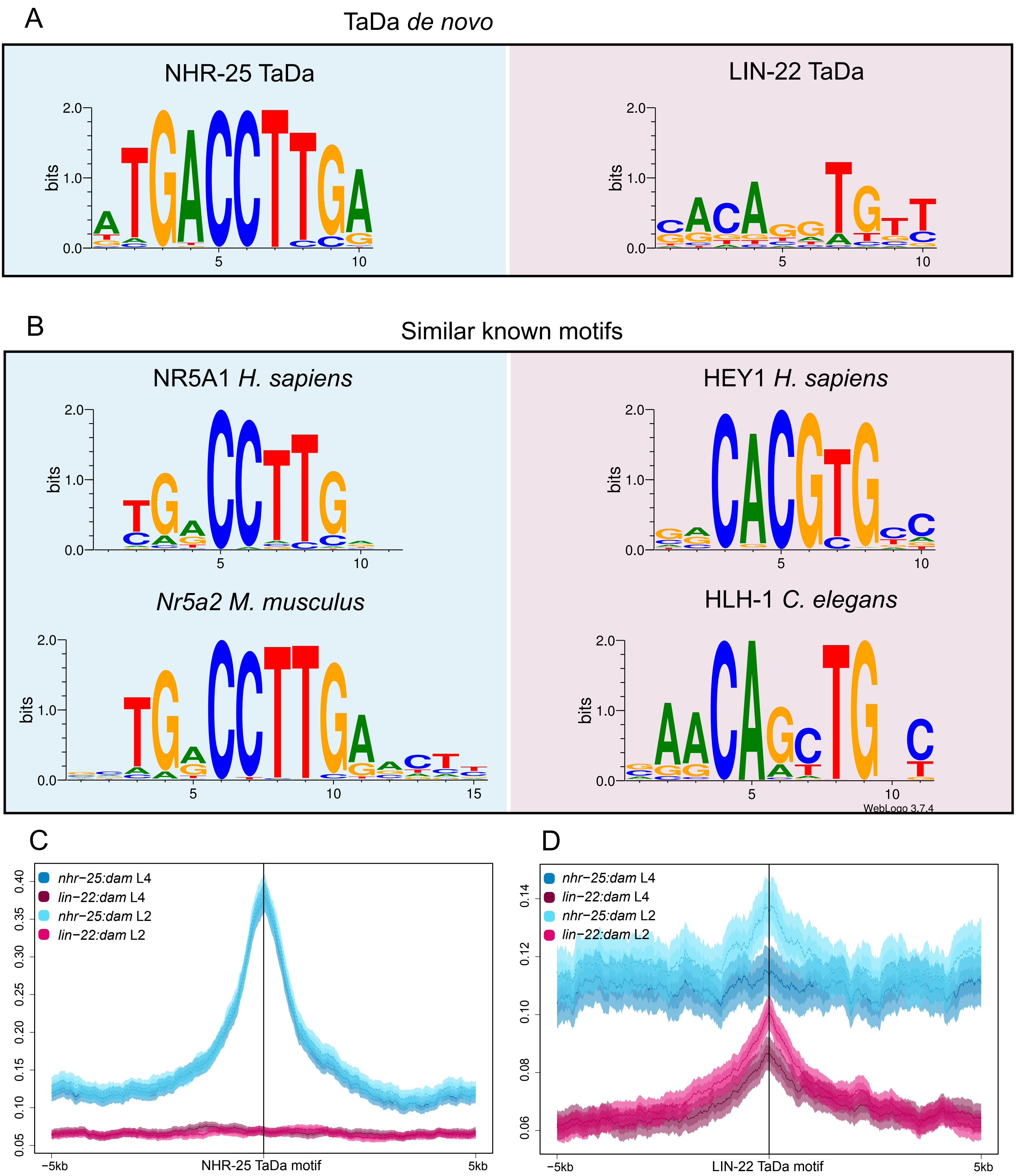

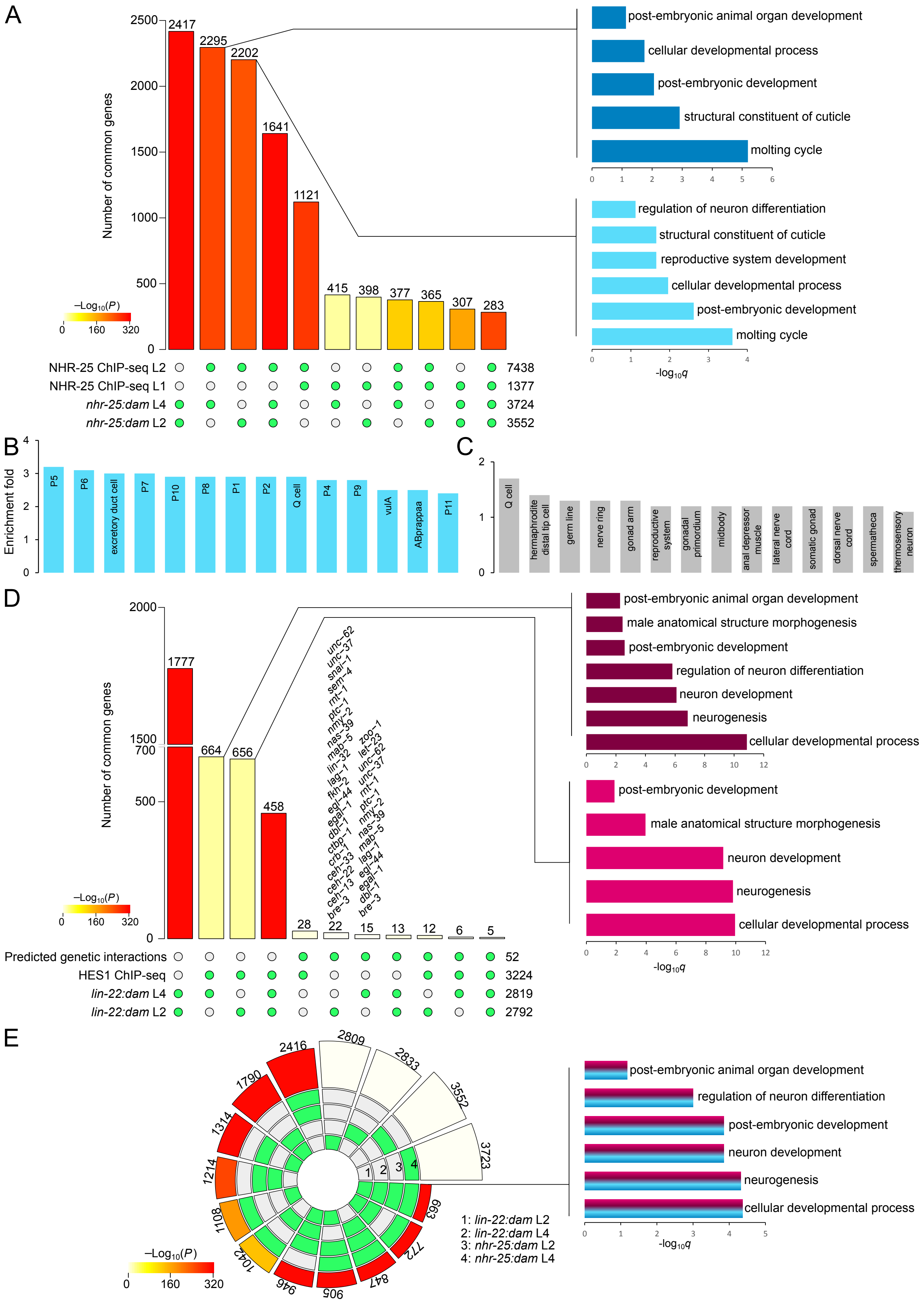

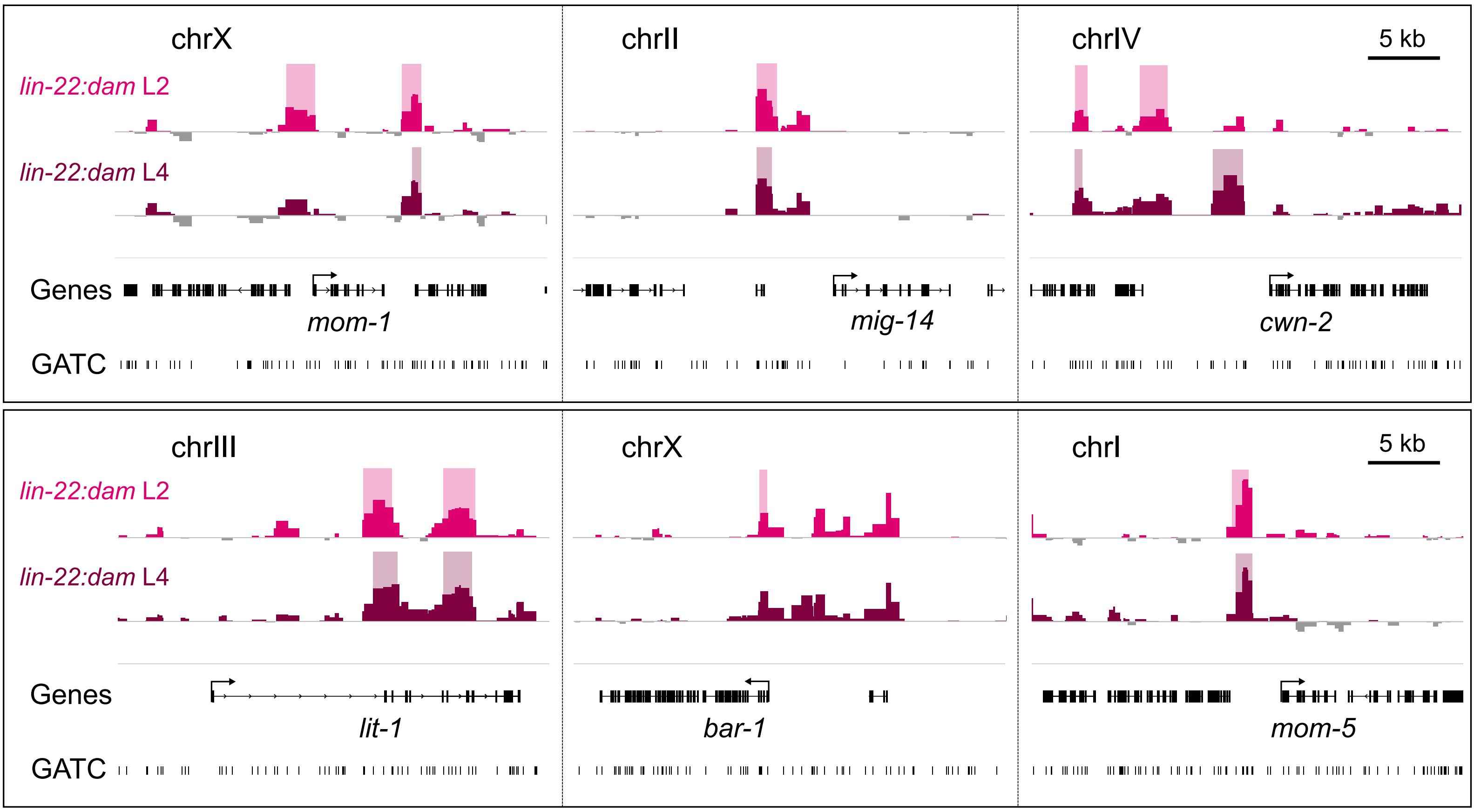
